# Heteromeric GABA_A_ receptor structures in positively-modulated active states

**DOI:** 10.1101/338343

**Authors:** Paul S. Miller, Simonas Masiulis, Tomas Malinauskas, Abhay Kotecha, Shanlin Rao, Sreenivas Chavali, Luigi De Colibus, Els Pardon, Saad Hannan, Suzanne Scott, Zhaoyang Sun, Brandon Frenz, Gianni Klesse, Sai Li, Jonathan M. Diprose, C. Alistair Siebert, Robert M. Esnouf, Frank DiMaio, Stephen J. Tucker, Trevor G. Smart, Jan Steyaert, M. Madan Babu, Mark S. P. Sansom, Juha T. Huiskonen, A. Radu Aricescu

## Abstract

Type-A γ-aminobutyric acid (GABA_A_) receptors are pentameric ligand-gated ion channels (pLGICs), typically consisting of α/β/γ subunit combinations. They are the principal mediators of inhibitory neurotransmission throughout the central nervous system and targets of major clinical drugs, such as benzodiazepines (BZDs) used to treat epilepsy, insomnia, anxiety, panic disorder and muscle spasm. However, the structures of heteromeric receptors and the molecular basis of BZD operation remain unknown. Here we report the cryo-EM structure of a human α1β3γ2 GABA_A_R in complex with GABA and a nanobody that acts as a novel positive allosteric modulator (PAM). The receptor subunits assume a unified quaternary activated conformation around an open pore. We also present crystal structures of engineered α5 and α5γ2 GABA_A_R constructs, revealing the interfacial site for allosteric modulation by BZDs, including the binding modes and the conformational impact of the potent anxiolytic and partial PAM, bretazenil, and the BZD antagonist, flumazenil. These findings provide the foundation for understanding the mechanistic basis of GABA_A_R activation.

---

The common architecture of eukaryotic pLGICs is now established, with at least one crystal or cryo-EM structure determined for the cation-selective nicotinic acetylcholine (nAChR) and serotonin type 3 receptors (5HT_3_R), and the anion-selective GABA_A_, glycine (GlyR), and GluCl receptors^1–9^. Each subunit extracellular domain (ECD) of 200-250 amino acids comprises an N-terminal α-helix and ten β-strands forming a twisted β-sandwich. Each TMD comprises an α-helical M1-M4 bundle, the M3 and M4 helices being connected by a large intracellular loop of 85-255 residues^10^. The two orthosteric neurotransmitter binding sites are situated between ECD principal (P) and complementary (C) faces, which intercalate around a water-filled vestibule that funnels into the ion channel transmembrane pore lined by five M2 helices. The structures have brought considerable understanding of the molecular basis of receptor function in homomeric receptor formats. However, heteromeric pLGIC structures have only been solved for nAChRs^1,2^. This is a key limitation given that the vast majority of mammalian pLGICs are heteromers possessing heightened complexity both molecularly and pharmacologically.

The GABA_A_R family comprises 19 different subunit subtypes: α1-6, β1-3, γ1-3, δ, ε, θ, π and ρ1-3^11^. Selective neuronal expression of particular subtypes, along with preferential assembly rules, ensure that the majority of GABA_A_Rs in the human brain comprise 2 a-subunits, 2 β-subunits and 1 γ-subunit, with a1, β2, β3, and γ2 subtypes exhibiting widespread overlapping expression profiles^12,13^. Alternative subtypes perform more specialised roles, for example, the α5 subtype influences cognition^14^ and moderation of α5β2/3γ2 receptors ameliorates animal model diseases of autism and Down syndrome, associated with cognitive deficits^15,16^. Modulation of α5β2/3γ2 receptors also improves recovery after stroke in a rodent model^17^. The homomeric GABA_A_R structures so far solved, GABA_A_R β3 and chimaeras that include *α* subunit TMDs, possess five-fold symmetry and will possess five identical copies of agonist and anaesthetic binding pockets^18,19^. In contrast, aβγ GABA_A_Rs possess two orthosteric GABA binding sites at β-a interfaces between the ECDs, two non-GABA binding a-β and γ-β interfaces, and one BZD binding site located at the a-γ interface^20^. Similarly, within the TMDs, aβγ GABA_A_Rs have at least three non-equivalent potential anaesthetic binding pockets. In the case of GABA_A_R β3, solved in a desensitized state, each subunit is bound by agonist and the subunit conformations are indistinguishable^5^. However, it is not known if different subunits within a heteromer adopt equivalent conformations in the activated state.

PAMs, such as BZDs, the intravenous general anaesthetics propofol and etomidate, barbiturates, endogenous neurosteroids, and alcohol (ethanol), bind GABA_A_Rs to promote their activation and channel opening, and thereby enhance GABAergic signalling^18–25^. This makes BZDs essential treatments for hyper-excitability disorders such as anxiety, insomnia and epilepsy^24^. However, these agents lack receptor subtype selectivity and cause unwanted sedation, addiction, and motor and cognitive impairment^26^. Agents with improved selectivity can ameliorate these side effects and allow for more selective targeting to treat other neurological disorders, including autism, Down syndrome, neuropathic pain, schizophrenia and stroke^15–17,27,28^. Structures of heteromeric GABA_A_Rs are essential to understand the molecular basis of inhibitory neurotransmission and drug subtype selectivity, and are anticipated to expedite the delivery of selective therapeutics against these disorders.

## RESULTS

### Structures of GABA_A_R constructs

Our initial goal was to obtain high resolution structures of the *α* and γ human GABA_A_R subunits. In addition to the β subunit structure we previously reported^5^, these are the essential constituents of the principal heteromeric receptor subtypes in the CNS^29^. Moreover, the α/γ interfaces harbour the BZD allosteric site^30–37^, which is of major pharmacological importance. Iterative rounds of screening and engineering of α5 subunits for pentamer monodispersity and yield identified a construct with 12 residue swaps from the β3 subunit, which readily forms homopentamers^5^, and 11 residue swaps from the BZD site complementary (C) face of the γ2 subunit. This construct recapitulated 100 % residue identity within the BZD site to the wild type α5-γ2 subunit interface (construct designated α5_HOM_; site designated BZD_HOM_; **Supplementary Fig. 1a-d**)^35–37^. Of particular note, an Asn114Gly substitution (β3 subunit glycine) to remove an N-linked glycosylation site that faces the extracellular vestibule and restricts homomeric assembly of α-subunits (observed in the cryo-EM construct – discussed below) was crucial to achieving monodispersity and pentamerisation (**Supplementary Fig. 1e**). We solved its structure to 2.6 Å resolution in complex with the drug, flumazenil^38^ (Anexate), a BZD site antagonist used to treat overdosage (**Fig. 1a, b** and **Table. 1**). Notably, although α5_HOM_ bound the BZD radioligands ^3^H-flunitrazepam and ^3^H-flumazenil, the affinity was 60-fold and 150-fold lower than for wild-type α5β3γ2 receptors (**Supplementary Fig. 2a-e**). To solve the structure of a BZD site with higher affinity we performed further screening and engineering on this background for heteromeric combinations containing γ2 subunits. These were transfected at 9:1 ratios of α5 to γ2 DNAs to favour one γ2 per pentamer. This identified a construct formed from α5_HOM_ subunits containing 4 additional residue swaps from the β3 subunit, and a chimera subunit comprising the γ2 ECD and α1 TMD, designated α5γ2_HET_. We crystallised this construct in complex with the BZD partial PAM, bretazenil^39^ and solved its structure to 2.5 Å resolution (**Fig. 1c, d; Table 1**). In contrast to α5_HOM_, inclusion of the chimeric γ2 subunit resulted in an additional higher apparent affinity site, designated BZD_HET_ (as well as the lower affinity BZD_HOM_ sites), which had only 2-fold, 5-fold and 15-fold lower affinity for ^3^H-flunitrazepam, ^3^H-flumazenil and bretazenil respectively, compared to wild-type receptors (**Supplementary Fig. 2a-f**).

**Figure 1.**
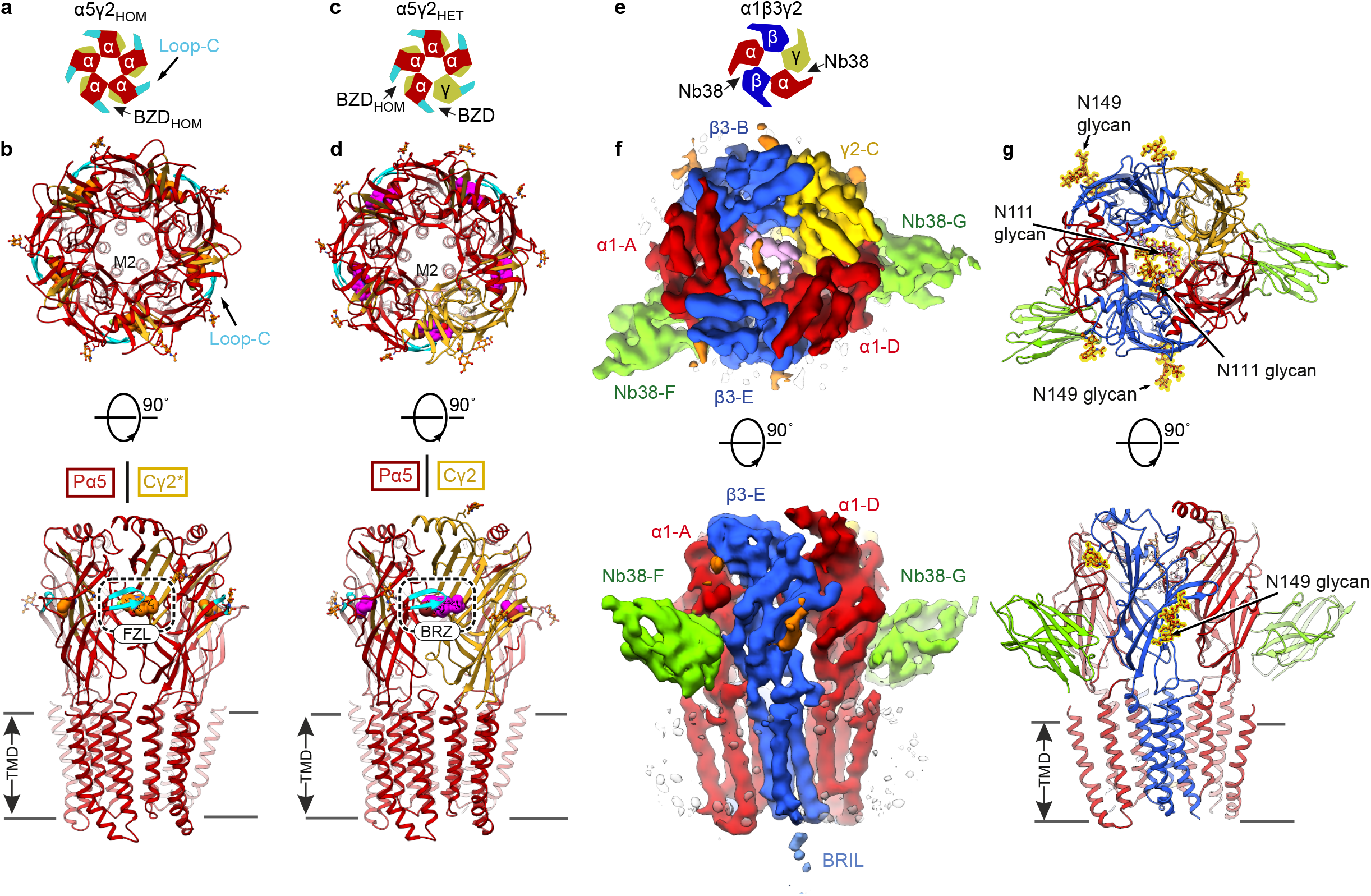
Architecture of GABA_A_Rs bound by BZDs and a Nb38 PAM. (**a**) Schematic top-down view of the subunit make-up of α5_HOM_. The BZD_HOM_ site is formed from homomeric α5 subunits (red) with the complementary (C)-face of the pocket engineered to contain γ2 residues (yellow). Loop C in cyan. (**b**) Crystal structure of α5_HOM_ pentamer shown in top-down and side-on views with flumazenil bound at BZDHOM sites shown with carbon atoms in orange (oxygen – red, nitrogen – blue, fluorine – cyan), inside dashed box for side view. A single M2 helix of the five pore-lining helices is labelled in the top-down view. N-linked glycans are shown as small orange spheres and sticks coloured by atom type (carbon atoms in orange, oxygen atoms in red). The principal (P) and complementary (C) faces between subunits in the side-on view are shown, boxed, for the BZD site between the foremost two subunits. (**c**) Schematic of α5γ2_HET_ showing a BZD_HOM_ site replaced by a BZD_HET_ site. (**d**) Equivalent views to (**b**) for α5γ2_HET_. Bretazenil shown with carbon atoms in magenta (oxygen – red, nitrogen – blue, bromine – brown). (**e**) Schematic of a1β3γ2_EM_ indicating interfaces bound by Nb38. (**f**) The 5.17 Å resolution cryo-EM map, top and side views. a1 subunits, red, β3 with thermostabilised apocytochrome b562RIL (BRIL) inserted in intracellular loop, blue, γ2, yellow, Nb38, green, detergent belt, white, N-linked glycans, orange except a1-A Asn111-linked glycan, pink. (**g**) a1β3γ2_EM_ atomic model. N-linked glycans are in ball-and-stick representation, outlined in yellow.

Attempts to crystallize tri-heteromeric GABAARs were unsuccessful, therefore we generated new constructs for single-particle cryo-electron microscopy. We designed α1 and γ2 subunit constructs in which only the M3-M4 intracellular loops were substituted by a short linker, SQPARAA^40^, and β3 subunits in which the M3-M4 intracellular loop was substituted by a thermostabilised apocytochrome b562RIL (BRIL)^41^ domain for unambiguous subunit identification. These constructs were co-expressed and assembled into functional α1β3γ2 GABA_A_Rs, designated α1β3γ2_EM_ (**Supplementary Fig. 3**). Only the γ2 subunit possessed a purification tag, in order to guarantee its presence in the pentamers. To facilitate particle alignment, we raised nanobodies against GABA_A_Rs and selected one of these, Nb38, that binds to a1 subunits. We obtained a 5.17 Å (FSC=0.143) cryo-EM map of a1β3γ2_EM_, in the presence of a saturating 1 mM concentration of GABA (**Fig. 1e-f and Supplementary Figs. 4 and 5; Table 2**). Two Nb38 molecules were bound between non-adjacent ECD interfaces, consisting of a1(P) faces and neighbouring β3(C) and γ2(C) faces, respectively, validating a1 subunit inclusion and receptor stoichiometry. At this resolution, the TMD a-helices of a1 and β3 subunits showed helical turns and side-chain densities for several large hydrophobic amino acids, although β-sheets were not fully separated into strands (**Supplementary Fig. 6**). The N-linked glycans served as markers that further allowed unambiguous subunit identification. Mannose branching could be clearly seen, for example, at a1 Asn111 and β3 Asn149. The detergent belt, BRILs and nanobody edges were largely disordered (**Supplementary Fig. 5g**). The γ2 subunit TMD was also less ordered, but otherwise the local resolution for the rest of the pentamer was consistent throughout (**Supplementary Figs. 5g and 6**). Viewed from above, the cryo-EM map confirms a clockwise subunit arrangement of a1-A, β3-B, γ2-C, a1-D, β3-E, consistent with previous indirect studies^42,43^. Despite the low resolution of this map, availability of high resolution crystal structures of individual subunits from α5γ2_HET_ and the previously solved GABA_A_R-β3_cryst_^5^ allowed us to build a model of the heteromeric α1β3γ2_EM_ receptor (**Fig. 1g; Table 2**). Inclusion of BRIL domains between the M3-M4 helices of the β3 subunit did not distort these helices (RMSD = 0.98 Å between a1β3γ2_EM_ β3 and GABA_A_R β3_cryst_ across 54 M3-M4 equivalent C_α_ positions).

### The BZD binding site and BZD binding modes

The ECD pockets between α5_HOM_ and α5γ2_HET_ subunits contained large positive peaks in the *F*o–*F*c electron density maps, which were unambiguously assigned to the co-crystallisation ligands flumazenil and bretazenil, respectively (**Supplementary Fig. 2i-l**). For α5γ2_HET_ the electron density map revealed inclusion of a single γ2 chimera subunit per pentamer, distinguishable by its unique side chains and glycosylation sites. This confirms the transfection ratio of 9α5:1γ2 ensured a pure stoichiometric population, as required to enable crystallization, and for radioligand binding studies. The lower apparent affinity of the BZDHOM sites did not affect the ligand binding mode which was the same as at the BZDHET site (**Supplementary Fig. 2m, n**). Furthermore, the binding modes of both flumazenil (observed only in BZD_HOM_ sites in α5_HOM_) and bretazenil are similar (**Figure 2a-h**). BZDs such as flunitrazepam (which lack the imidazo C(3)-linked ester moiety and instead possess a diazepine C(6) phenyl ring) exhibit a distinct pharmacology from flumazenil and bretazenil because they require a histidine residue in the BZD site of α1/2/3/5 subtypes for binding^32,44^. Radioligand binding on a modified α5γ2_HET_ construct, α5γ2_HETΔ_, with BZD_HOM_ sites removed (by reversing the γ2 substitutions in the α5 subunits), so that only the BZDHET site remained, revealed that a His105Arg substitution ablated ^3^H-flunitrazepam binding but retained the same apparent affinity for flumazenil (**Supplementary Fig. 2g, h**). Thus, the differential impact of the His to Arg substitution on the binding of distinct classes of BZDs to wild type receptors is reproduced in this construct.

**Figure 2.**
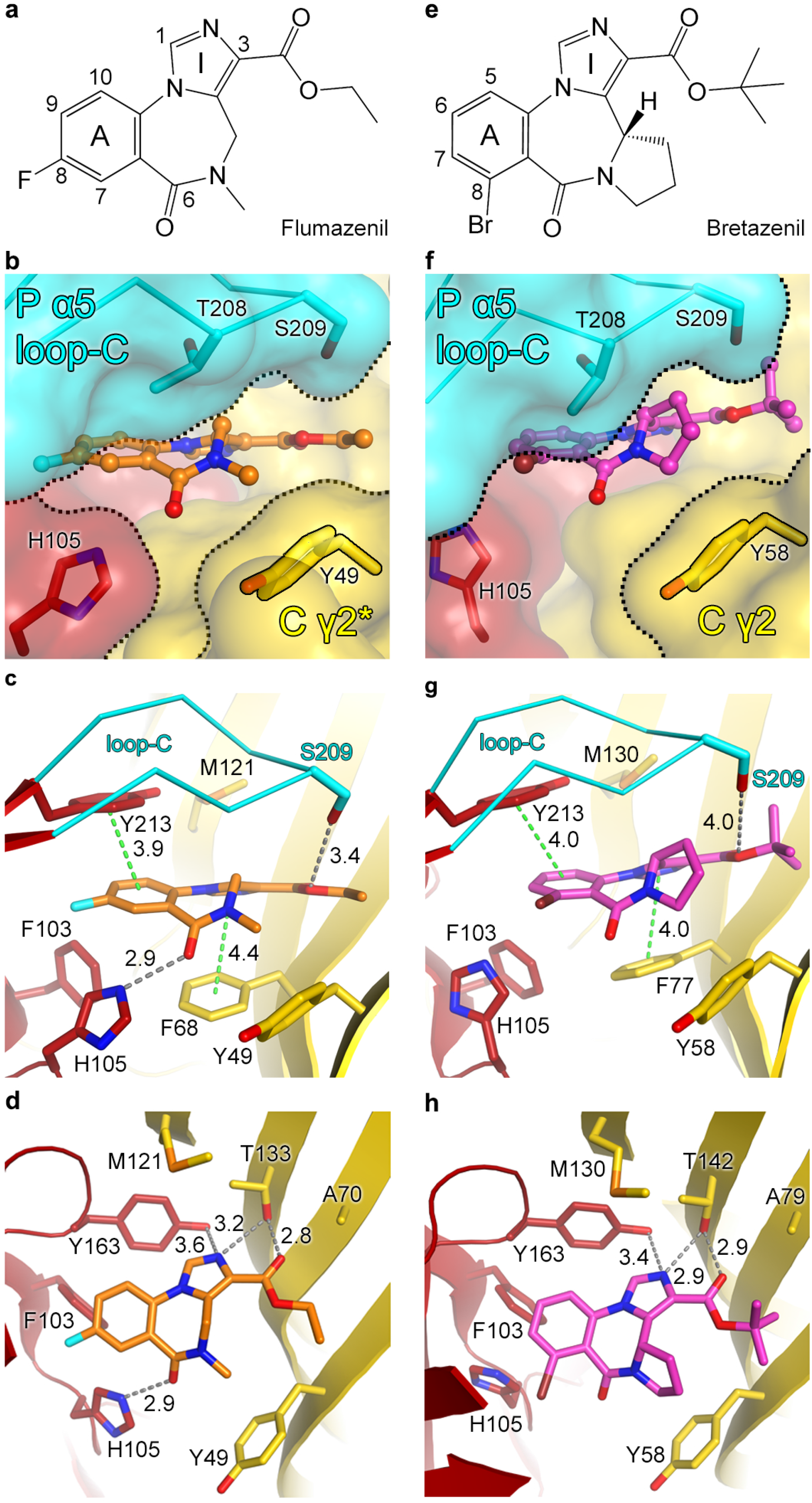
Binding modes of bretazenil and flumazenil at the BZD_HET_ and BZD_HOM_ sites. (**a**) Structural formula of flumazenil. Carbon atoms referred to in the text are numbered. Imidazole (I) and benzene rings (A) are labelled. (**b**) Surface representation of the BZD_HOM_ binding pocket (a5 subunit, red; loop-C of a5 subunit, cyan; the complementary (C)-face containing γ2 residues, yellow, designated γ2*) with flumazenil shown as sticks (carbon, orange; oxygen, red; nitrogen, blue; fluorine, cyan). (**c**), (**d**) Side and top views of flumazenil bound to this BZD site. **e**, Structural formula of (S)-bretazenil. (**f-h**), Bretazenil bound to the BZD_HET_ site (carbon, magenta; oxygen, red; nitrogen, blue; bromine, brown). γ2 subunit, yellow. Side chains of amino acid residues contacting the ligand are numbered and shown as sticks. Distances (Å) between selected atoms and aromatic rings are shown as grey and green dashed lines, respectively.

Intermolecular binding contacts between the ligands and receptor occur predominantly through van der Waals (vdW) interactions, which span the interface between subunits. From the α-subunit (P)-face, five hydrophobic residues make vdW contacts with the ligand benzene A-rings: Phe103 and His105 from the β4 strand (historically named loop A^10^); Tyr163 from the β7-β8 loop (loop B); Ile206 and Tyr213 from the β9-10 hairpin (loop C) (**Fig. 2 a-h and Supplementary Fig. 1**). Of particular importance are Tyr163, which forms T-shaped π-stacking interactions with the A and I ligand rings, and Tyr213, forming parallel π-stacking interactions with the A rings. Consistent with such interactions, Tyr163Phe and Tyr213Phe substitutions retain BZD potentiation whilst Ser substitutions reduce sensitivity at least 10-fold^33^. The structures also explain why moving the flumazenil C(8) fluorine to C(9) or C(10) chlorine or adding a C(1) methyl substituent (Compounds 1-3 in **Supplementary Fig. 7a-d**) reduce binding by over 100-fold^45–47^. These substituents clash with loop B Tyr163, as shown by *in silico* docking studies (**Supplementary Fig. 7a-d**). The lowest free energy binding mode of flumazenil overlays the one observed experimentally in a5_HOM_, whereas compounds 1-3 are displaced by ∼2 Å away from Tyr163, impairing the T-shaped π-stacking interactions. In contrast, the C8 azide derivative Ro15-4513^48^, a competitive BZD antagonist developed as an antidote to alcohol and more recently used as a PET ligand^49^, docked similarly to flumazenil, with the azide positioned under the loop C Tyr213, which it chemically photolabels^50^ (**Supplementary Fig. 7e**).

The loop-A His105 undertakes a ligand-dependent reorientation within the site (**Fig. 2b-d, f-h**), its side chain rotating about the C_α_-C_β_ bond to accommodate either the bretazenil bromine or the flumazenil chlorine atoms (at adjacent positions on the A rings, **Fig. 2a, e**). As shown by the radioligand data (**Supplementary Fig. 6g, h**), a His105Arg substitution does not affect flumazenil binding even though it ablates flunitrazepam binding. The explanation for this is observed in the γP/αC site in α5γ2_HET_, where the equivalent residue to His105 is γ2 Arg114, which is well accommodated under the bretazenil ligand (**Supplementary Fig. 7i, j**).

From the (C)-face, the β2 strand (loop D) phenylalanine side chain (γ2 Phe77 in the BZD_HET_ site, γ2* Phe68 in the BZD_HOM_ site) forms π-stacking interactions with the ligand I rings (**Fig. 2c, g**). These explain why a Phe77Ile substitution reduces flumazenil affinity 1000-fold and why GABA_A_ receptors containing the γ1 subunit, where an Ile residue occupies the equivalent position, are much less responsive to BZDs^36^. Above Phe77, an alanine residue (γ2 Ala79 in the BZD_HET_ site, γ2* Ala70 in the BZD_HOM_ site) demarcates the top of the BZD site and faces the I ring C(3) substituent, consistent with a previous mutagenesis study predicting close apposition of these two elements^51^ (**Fig. 2d, h**). The neighbouring β6 strand (loop E) threonine (γ2 Thr142 in the BZD_HET_ site, γ2* Thr133 in the BZD_HOM_ site) forms putative hydrogen bonds with both the imidazole nitrogen and ester carbonyl of the ligands, consistent with original pharmacophore models proposing a dual H-bond contribution from these moities^52^ (**Fig. 2d, h**). The ester carbonyl group is essential for flumazenil binding versus ketone or ether functionalities^46^, whilst other derivatives that maintain the isosteric constraints retain high affinity binding^52,53^.

Patients on BZDs can experience a variety of adverse events, including cognitive and psychomotor effects, tolerance and, in some cases, paradoxical behaviours such as disinhibition leading to aggression^26^. To explore whether there is genetic variation within the BZD site, which may contribute to response variability in individuals, we mapped allelic variants of 138,632 unrelated healthy humans (gnomAD database^54^) for α1-6 and γ1-3 subunit residues within 5Å of ligand bound to α5γ2_HET_ and α5_HOM_ (**Fig. 3a-c**). Although no common allelic variants were found, many rare allelic variants (ranging from 1 in 1,000 to 1 in 100,000 people) were identified, spanning the four major physiological BZD site subtypes, α1γ2, α2γ2, α3γ2 and α5γ2. The α5 subunit Gly161Val African variant is predicted to sterically disrupt the folding of loop B reducing ligand-binding affinity, as shown previously for the orthosteric GABA site reducing agonist binding 400-fold^55^. Several African, East Asian and European Non-Finnish variants cluster at loop A His105, including α1 subunit tyrosine or arginine substitutions and α3 subunit tyrosine or asparagine substitutions, all of which have been shown to reduce sensitivity to classical benzodiazepines 10-20-fold^56^. Finally, the loop C tip (Ser209-Thr210-Gly211 in the α5 subunit) experiences the highest incidence of variation in all four BDZ binding α1/2/3/5 subunits. Substitutions within this region have previously been shown to reduce sensitivity to potentiation by classical BZDs and the sedative zolpidem, a non-BZD that binds the BZD site, by 5-10-fold^34,57^. Whilst it might be expected that reduced binding will result in a suboptimal patient response to BZDs, for some of these allelic variants the responses to anxiety or epilepsy treatment might actually be improved. The GABA_A_R α1 subunit is linked to unwanted sedative and addictive side effects^58^ so individuals possessing α1 His102 substitutions might, intriguingly, experience less side effects.

**Figure 3.**
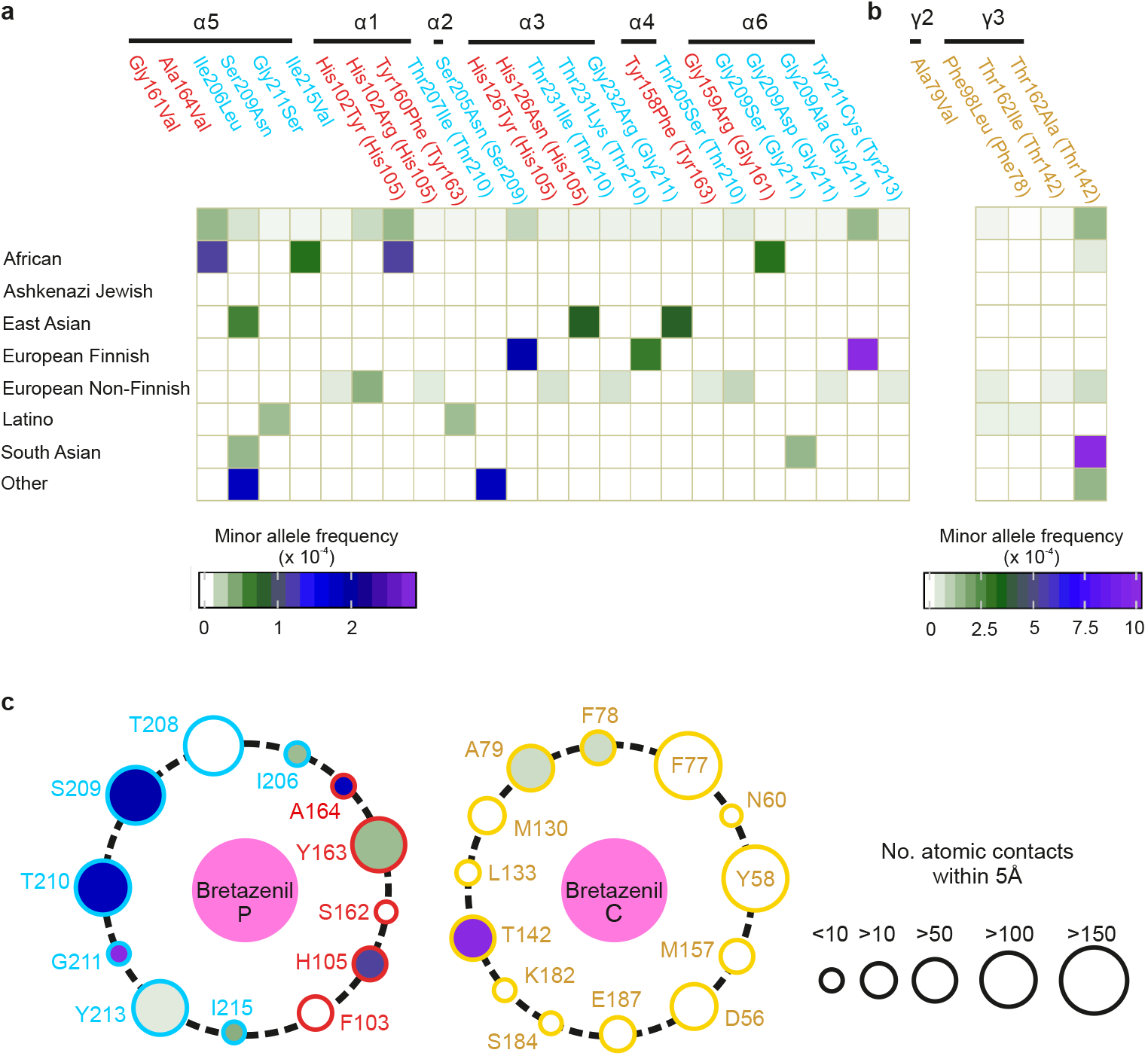
Sequence variability at the BZD binding pocket in the human population. (**a**) and (**b**) Heat maps of allelic variants from GABA_A_R α and γ subunits respectively, for residues within 5Å of bound BZD in a5γ2_HET_ and a5_HOM_, across ethnic groups from the gnomAD database^54^ of 138,632 unrelated healthy individuals. Boxes are colour coded by frequency, with intensity indicators underneath. Text colour corresponds to residue location: red – α-subunit loops A or B; blue – α-subunit loop C; yellow – γ-subunit. Residue numberings are for mature sequences (Uniprot: α1 P14867, α2 P47869, α3 P34903, α4 P48169, α5 P31644, α1 Q16455, γ2 P18507, γ3 Q99928) with α5γ2_HET_ numbering in parentheses where appropriate. (**c**) Asteroid plots of ligand-residue contacts between bretazenil and α5 P-face and γ2 C-face residues. Circle size reflects contact number (size indicator provided). Circle outline colour corresponds to residue location (same as for text in, **a**). Circles are filled for residues with allelic variants, and coloured according to frequency (same as in, **a**).

### Nb38 is a novel positive allosteric modulator

While the cryo-EM map resolution limits our ability to define precise side-chain interactions, it is clear that major receptor-Nb38 contact points exist between: the CDR2/3 loops and β9/β10 strands and β9-β 10 hairpin (loop C), with CDR2 inserting into the pocket under loop C; CDR1-3 loops and the β6/β7 strands and β6-β7 loop (Cys-loop); CDR1/2 loops and the (C)-face β1/β2 strands and β8/β9 loop (loop F) (**Fig. 4a, b** and **Supplementary Fig. 8a, b**). Nb38 binding affinity (K_D_) for detergent-solubilised α1β3γ2_EM_ receptors, as determined by surface plasmon resonance (SPR), was increased 6.5-fold in the presence of 1 mM GABA, from 1.61 nM to 248 pM, suggesting that it favours binding and stabilises an activated receptor conformation (**Supplementary Fig. 8c, d**). Whole cell patch-clamp recording confirmed this. 10 μM Nb38 strongly potentiated EC_10_ GABA currents by 480 ± 30 *% (n* = 7) for α1β3γ2_EM_ and 290 ± 20 *% (n* = 7) for wild-type (WT) α1β3γ2 (**Fig. 4c**), greater than achieved by the BZD diazepam, 180 ± 20 % (n = 12) for α1β3γ2_EM_ and 130 ± 20 % (n = 9) for α1β3γ2_WT_ (**Supplementary Fig. 3b**). Application of Nb38 alone at concentrations up to 10 μM had only a weak direct agonist effect (**Fig. 4c**). Importantly, Nb38 was α1 subunit selective, eliciting no potentiation of wild-type a2-a6β3γ2 receptors (**Fig. 4d**). Thus, Nb38 is a novel pharmacological tool with superior efficacy and selectivity for a1 subunit receptors over benzodiazepines. The Nb38 ability to potentiate GABA_A_Rs was also observed for spontaneous inhibitory post-synaptic currents (sIPSCs) from dentate gyrus granule cells (DGGCs) in acute hippocampal slices, which primarily stem from a1-subunit containing receptors^59^. Without significantly affecting sIPSC frequency or rise-time, 2 μM Nb38 increased amplitude 37 ± 9 % (*p* < 0.05) and prolonged decay times by 96 ± 19 % *(p* < 0.05) to increase GABAergic drive (in terms of charge transfer calculated as the average sIPSC surface area) by 94 ± 14 %, *n* = 4 *(p* < 0.01) (**Fig. 4e, f**). By comparison, 500 nM diazepam also significantly prolonged sIPSC decay time, although to a lesser extent (39 ± 10 %, *p* < 0.01 %), and did not significantly increase GABAergic drive.

**Figure 4.**
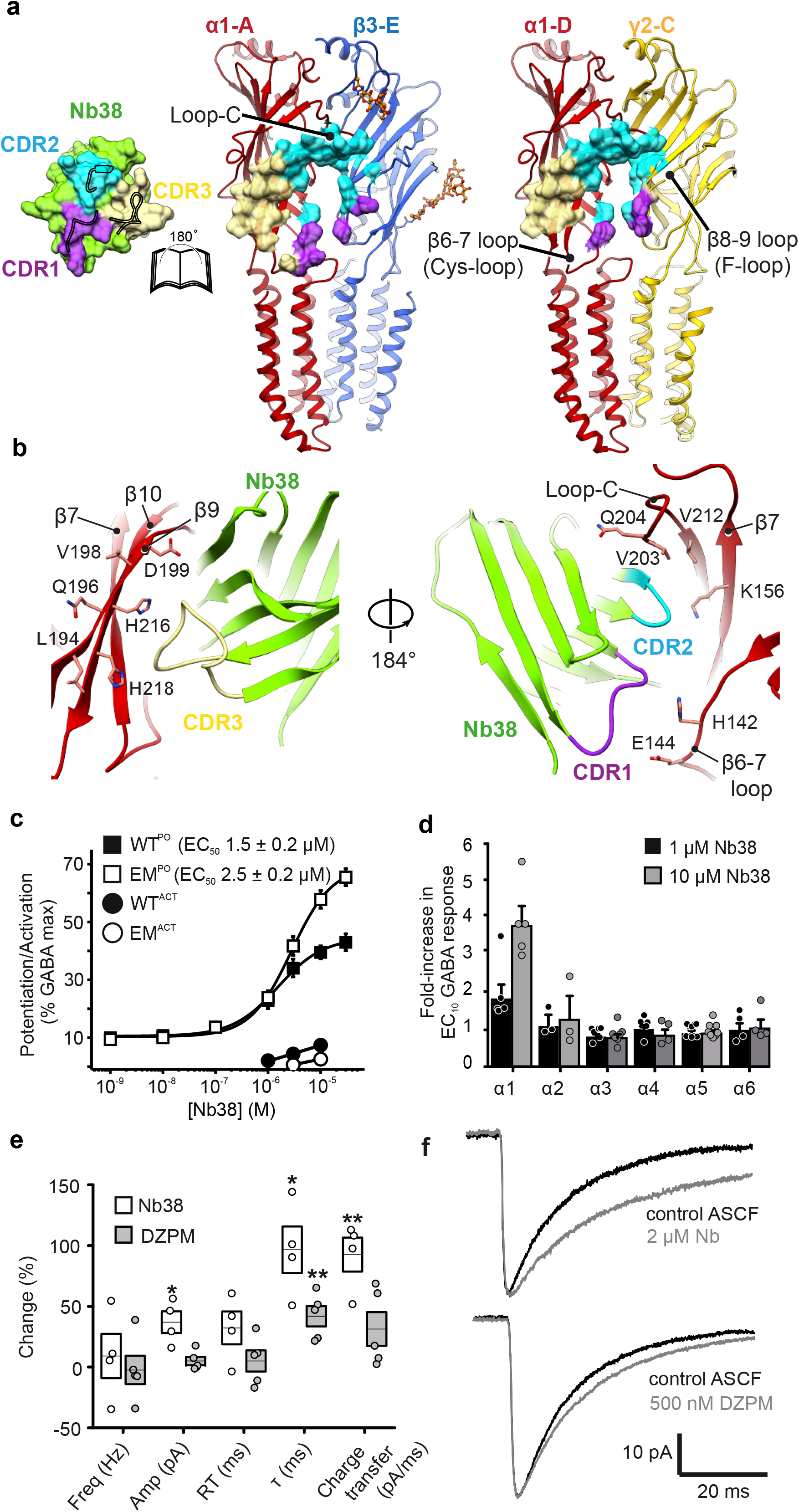
Nb38 binding and modulation of α1β3γ2 receptors. (**a**) ‘Open book’ view of the Nb38 interaction surface with α1β3γ2_EM_ at the α1-A-β3-E and α1-D-γ2-C (just under the BZD site) interfaces. Contact regions for: CDR1, purple; CDR2, cyan; CDR3, yellow. (**b**) Detailed view of Nb38 binding to the α1 subunit. CDR3 loop binding to β9 and β10 strands and alternate view rotated 184° degrees around the receptor longitudinal axis to show CDR1 and CDR2 interactions with the Cys-loop, loop-C, and β7 strand. Amino acids in α1 subunit which are predicted to contribute to Nb38 binding are highlighted. (**c**) Nb38 concentration-response curves for potentiation of GABA EC_10_ responses (WT^PO^, EM^PO^) and direct activation (WT^ACT^, EM^ACT^), respectively, measured from whole-cell patch-clamp recordings of HEK293T cells expressing either wild-type (WT) α1β3γ2 GABA_A_Rs or α1β3γ2_EM_. *n* = 7, EC_50_s are accrued from different cells. (**d**) Bar chart demonstrating α1-subunit selectivity of Nb38 compared to αnβ3γ2 GABA_A_Rs (n = 2-6). Note, no potentiation at other subtypes. *n* = 3-4, each data point comes from different cells. (**e**) Effect of Nb38 (2 μM) and diazepam (DZPM, 500 nM) relative to control ACSF on sIPSC frequency (freq) in dentate gyrus granule cells, amplitude (amp), rise-time (RT), decay time (τ) and charge transfer. \**p* < 0.05 and ** *p* < 0.01 significance compared to ACSF (paired t-test). Nb38 and DZPM were tested on 4 and 5 cells respectively. (**f**) Average sIPSC waveforms after incubation with Nb38 and DZPM (grey) superimposed over control ACSF (black) before incubation with ligand.

### Impact of N-linked glycosylation on pentamerization

The a1β3γ2_EM_ cryo-EM map revealed multiple N-linked glycans attached to the a1 (Asn111, β5-β5’ loop), β3 (Asn80, β3-strand and Asn149, β7-strand) and γ2 (Asn208, β9-strand) subunits (**Fig. 1c**). Of particular note, the two a1 Asn111 glycans occupy the ECD vestibule and, unexpectedly, adopt well-ordered conformations (**Fig. 5a**). The a1-D glycan projects upwards into the extracellular space (**Fig. 5b**). In contrast, the a1-A glycan projects horizontally to form putative CH-π interactions between the pyranose ring of a mannose moiety and the apposing Trp123 side chain from the γ2 β5-β5’ loop (**Fig. 5c, d**). This glycosylation site is conserved across all a-subunits (a1-6) and Trp123 is conserved across all γ-subunits (γ1-3), but not β-subunits (**Fig. 5e**). Thus, this glycan pairing is expected to exist in all aβγ GABA_A_Rs in the human brain, a feature absent from all previously solved pLGIC structures^10^. Enzymatic glycosylation of a1 Asn111 precedes exit from the endoplasmic reticulum (ER)^60^ and requires access to the inner face of a1, meaning that it must precede assembly and closure of the pentameric ring. The non-random orientation induced by the α1-A glycan interaction with γ2 Trp123 suggests that pentamerization comes from pre-assembled αβγ trimers with a horizontal glycan, followed by the addition of a second αβ dimeric unit (**Fig. 5f**). Furthermore, the Asn111 glycosylation represents a stoichiometric control mechanism, preventing the inclusion of more than two α-subunits via steric hindrance. Accordingly, we could only obtain the pentameric α5_HOM_ and α5γ2_HET_ constructs described earlier after mutating the Asn111 residue (**Supplementary Fig. 1e**).

**Figure 5.**
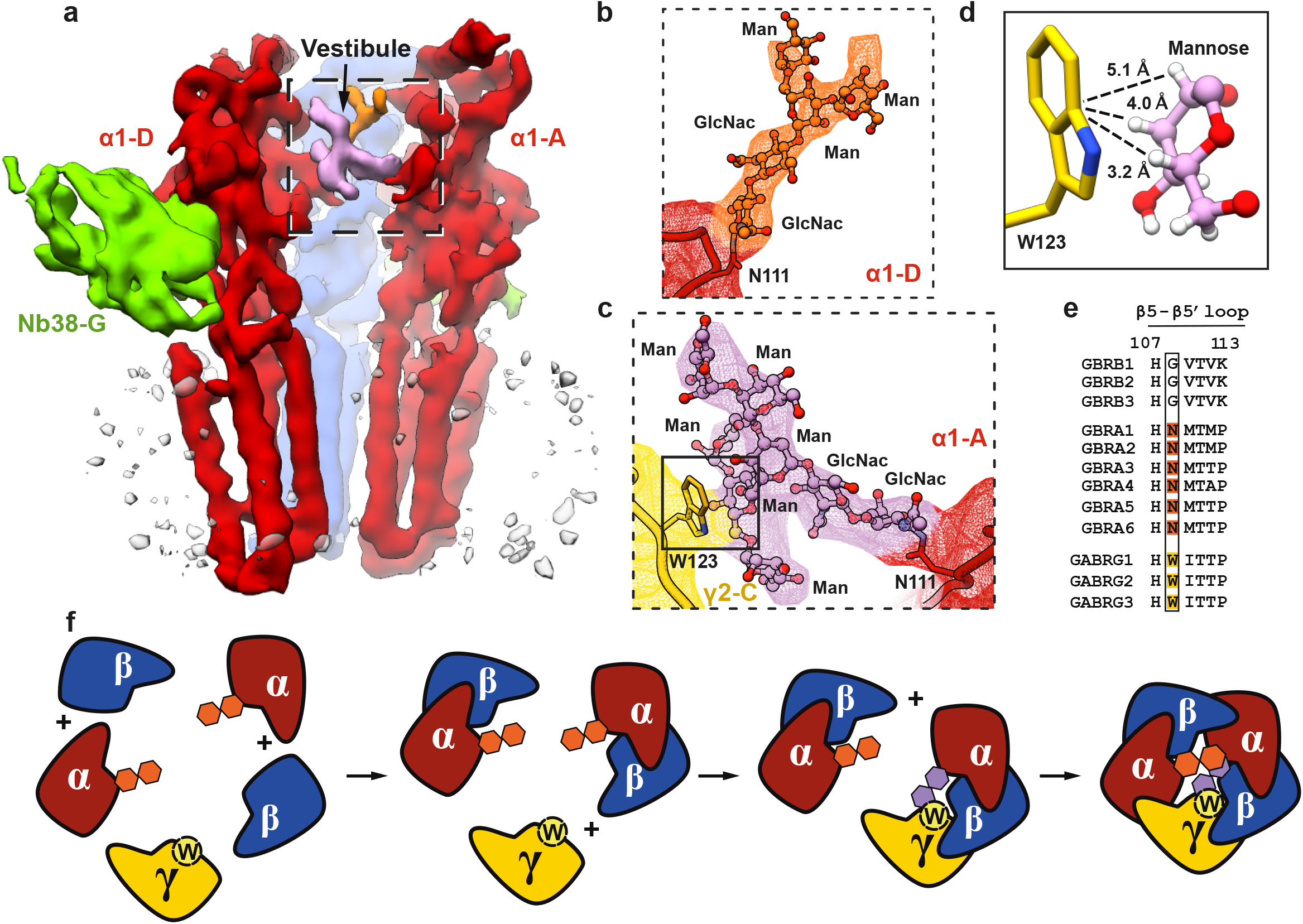
N-linked glycosylation of a1β3γ2_EM_ α1 subunits. (**a**) Side view of a1β3γ2_EM_ map showing a1 N-linked glycans filling the channel vestibule. β3-B and γ2-C subunits were hidden to visualise the vestibule. (**b**) and (**c**) Close-ups of a1-D and a1-A Asn111-linked glycans. Atomic model and cryo-EM maps are shown in orange and pink, respectively. (**d**) Putative CH-π interaction between the Trp123 side-chain and the pyranose ring of a mannose moiety from the a1-A Asn111-linked glycan. Distances between the centre of mass of the Trp123 indole and hydrogens of the interacting mannose residue are shown. (**e**) Sequence alignment of β5-β5’ loops. All a subunits contain N-linked glycosylation sites, and all γ subunits contain Trp residues at the equivalent position (boxed). Sequence numbering shown is based on β3 subunit (mature isoform 1). (**f**) Proposed assembly sequence for aβγ GABA_A_Rs. a1 and β3 subunits produced in the ER first form dimers through the β3(P)-a1(C) interface. An a1β3 dimer binds to the γ2 subunit through the γ2(P)-β3(C) interface with the Asn111-linked glycan on the a1 subunit interacting with γ2 Trp123 to stabilise the horizontal glycan conformer. Pentamerization occurs through β3(P)-a1(C) and a1(P)-γ2(C) interfaces between an a1β3γ2 trimer and an a1β3 dimer in which its Asn111-linked glycan adopts an upward trajectory.

### The extracellular region conformations

Superpositions of α5_HOM_, α5γ2_HET_ and GABA_A_R-β3_cryst_^5^ single ECDs, reveal that all three subunits comprising the major synaptic heteromer format (2α_n_-2β_n_-1γ_n_) adopt highly similar β-sandwich arrangements (RMSD in the 0.59-0.76 Å range between different chains over 208 equivalent Cα positions), with a notable divergence in the flexible β8-β8’ and β8’-β9 loops (loop F) (**Supplementary Fig. 8e**). Global superposition of the α5_HOM_, α5γ2_HET_ and α1β3γ2_EM_ ECD pentamers reveal that all the subunits have adopted positions relative to the pore axis similar to agonist-bound GABA_A_R-β3_cryst_ and the homologous agonist-bound glycine receptor^6^ (GlyR), rather than antagonist-bound GlyR (RMSD in the 0.7-1 Å range versus GABA_A_R-β3; 1.0-1.4 Å versus agonist-bound GlyR; 2.1-2.3 Å versus antagonist-bound GlyRs, over 208 equivalent Cα positions) (**Supplementary Fig. 8f-j**). The activated ECD bases swing out from the pore allowing it to widen while the loop C tips close inwards. Of note, the same assumed ECD positions between subunits means the α1β3γ2_EM_ BZD site is highly similar to the BZD sites of α5_HOM_ and α5γ2_HET_ (**Fig. 6a-c**). Consistent with this, *in silico* docked flumazenil and bretazenil assumed the same binding modes in α1β3γ2_EM_ (**Fig. 6d-e**). In the case of α5_HOM_ and α5γ2_HET_ this means the polar Thr208-Ser209-Thr210 residues at the tip of loop C sterically trap BZD in the site (**Fig. 2b, f**). The tight closure is incompatible with the inverted concave tetracyclic ring series of the (*R*)-bretazenil enantiomer^47^, as confirmed by *in silico* docking in which the lowest energy binding mode of (*S*)-bretazenil recaptures the binding mode in a5γ2_H_E_T_ whereas (*R*)-bretazenil is pushed out of the pocket (**Supplementary Fig. 7f, g**). The activated ECD conformations of a5γ2_H_E_T_ and a1β3γ2_EM_ are consistent with them being bound by PAMs, i.e. bretazenil and Nb38, respectively. That all five subunits of a1β3γ2_EM_ adopt equivalent ECD conformations despite non-equivalent occupancy of the inter-subunit pockets (two β3P/a1C orthosteric sites presumed to be bound by GABA; one a1P/β3C and one a1P/γ2C site bound by Nb38; one a1P/β3C site presumed empty) holds to the Monod-Wyman-Changeux (MWC) theory that postulates transitions between states (e.g. closed-to-open states) preserve symmetry of the oligomer^61^. In the case of a5_HOM_, the bound ligand, flumazenil, is a neutral BZD antagonist with no preference between activated or resting states^53^, which will bind either equally well. In this instance a5_HOM_ has adopted the activated conformation as is the case for a5γ2_HET_, which might reflect an intrinsic preference of the construct ECDs to favour the activated conformation, as previously observed for β3 ECDs in the absence of bound ligand^62^. Interestingly, increasing the size of the C(3)-linked ester substituent of flumazenil analogues correlates with PAM activity^53^. A flumazenil analogue where the ester is substituted by an alkyne of the same length was a neutral modulator (antagonist), whilst addition of a *tert-* butyl group, as is present in bretazenil, conferred PAM activity^46^. *In silico* docking of this PAM analogue (compound 4) into a5γ2_HET_ resulted in the lowest free energy binding mode directly overlapping with bretazenil (**Supplementary Fig. 7h**). In both cases, the *tert*-butyl groups are precisely positioned to bridge the gap between loop C Thr208-Ser209 and the adjacent subunit at β1-strand Asp56-Tyr58 (**Fig. 2b and Supplementary Fig. 7h**). Closure of loop C is primarily a consequence of the quaternary motion undertaken by the pLGIC ECDs during activation^6,8,10,20^. This suggests that, by bridging the subunits at this critical juncture, bretazenil and other BZDs with appropriately large C(3) substituents stabilise the integrally linked process of loop C closure and ECD quaternary activation via their larger contact surfaces compared to flumazenil.

**Figure 6.**
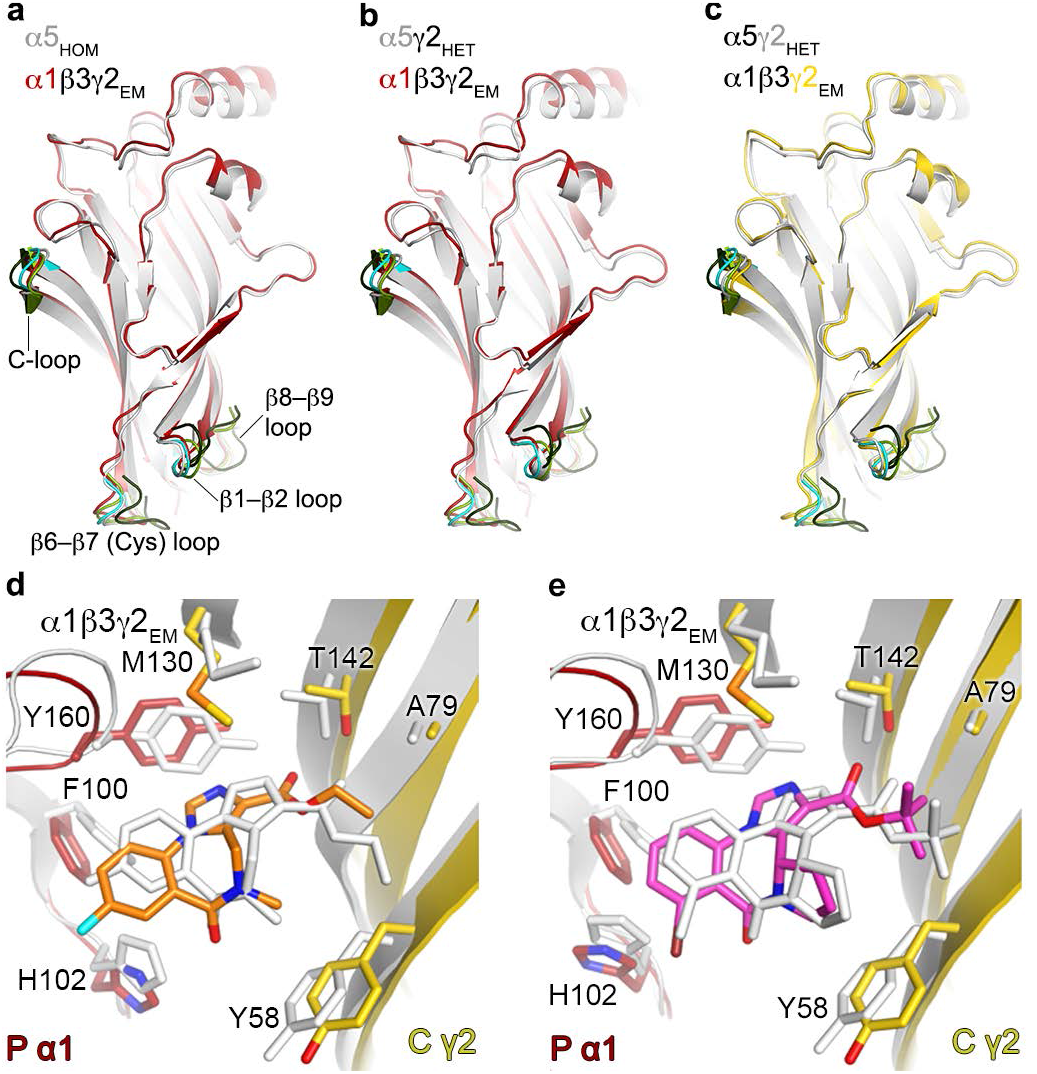
α5_HOM_, α5γ2_HET_ and α1β3γ2_EM_ ECD activated conformations and BZD binding modes. Side-on views of single ECDs from globally superposed pentamers comparing quaternary conformations relative to the pore axis (not shown – right of ECDs), shown looking into loop C. (**a**) α5_HOM_ (light grey) versus α1β3γ2_EM_ α1 subunit, (**b**) α5γ2_HET_ α5 subunit (light grey) versus α1β3γ2_EM_ α1 subunit, (**c**) α5γ2_HET_ γ2 subunit (light grey) versus α1β3γ2_EM_ γ2 subunit. Agonist binding C-loop and the ECD base loops (β1-2 loop, β6-7 loop – Cys-loop, β8-9 loop) are also shown for activated β3_cryst_ (PDB IDs: 4COF – cyan), agonist-bound (activated) glycine receptor (PDB ID: 5JAE – light green) and resting closed GlyR (PDB ID: 5JAD – dark green). The α1β3γ2_EM_ α1 and γ2 ECDs, and α5_HOM_ and α5γ2_HET_ α5 or γ2 subunits all occupy the activated conformation with loop-C closed and base loops tilted away from the pore axis, similar to previously solved activated β3_cryst_ and GlyR structures, and different from resting state GlyR. Computational docking of flumazenil (**d**, estimated free energy of binding, −8.9 kcal/mol) and bretazenil (**e**, −7.8 kcal/mol) to α1β3γ2_EM_. Docked ligands, α1 and γ2 subunits from α1β3γ2_EM_ are colored, crystal structures of GABA_A_R in complex with flumazenil (**d**) and bretazenil (**e**) are shown in light grey.

### Ion permeation pathway and pore conformation

α1β3γ2_EM_ possesses two positively charged rings, both previously observed in GABA_A_R-β3_cryst_, halfway down the vestibule and at the intracellular end of the pore (**Supplementary Fig. 9a-c**), and previously proposed as ion selectivity filters in pLGICs^63–65^. In the crowded environment of the glycan-filled vestibule, the ion permeation pathway is restricted to ∼ 5Å diameter (**Fig. 7a**), less than a fully hydrated Cl^−^ ion (6.1 Å)^66^ but larger than a dehydrated Cl^−^ ion (Pauling radius of 1.8 Å). However, the polar, semi-flexible glycan chains are not expected to impede the Cl^−^ flux significantly. The membrane spanning pore, lined by five M2 helices, one from each subunit, narrows from greater than 8 Å diameter on the extracellular side (excluding the uppermost concentric residue ring, designated 20’, which comprises flexible polar side chains that will not impede conductance), to ∼5.6 Å on the intracellular side, at the - 2’ ring (α1 Pro253, β3 Ala248, γ2 Pro263; **Fig. 7b**). Although slightly narrower than a fully hydrated Cl^−^ ion, this is wider than the closed −2’ desensitization gate of GABA_A_R-β3_cryst_^5,67^ (3.2 Å), and also dehydrated Cl^−^ ions, and is consistent with an open-channel state, within the 5.5-6 Å range previously estimated on the basis of different-size anion permeability^68^. It is important to state that although the M2 helices (lining the pore) of α1 and β3 subunits are well ordered in the cryo-EM map, the γ2 subunit TMD region is mobile. This appears to be an artefact caused by detergent solubilisation, as we observed a reduction of this motion upon addition of lipids (**Supplementary Fig. 4a**). Iterative rounds of particle classification led to a map in which the γ2 subunit TMD model could be unambiguously placed, but not accurately refined. Therefore, to ascertain the conductance state, we performed 100 ns molecular dynamics simulations^69^ on the TMD pore embedded in a POPC membrane with water and 0.15 M NaCl on each side and a + 0.12 V transmembrane potential (where the sign of the voltage refers to that of the cytoplasmic face). Water molecules occupied the length of the pore, indicating there was no significant hydrophobic barrier leading to de-wetting of the channel^70^. There was an influx of multiple Cl^−^ ions, equivalent to an estimated conductance of the order of 100 pS, sufficient to account for physiological conductances (25-28 pS^71^) (**Fig. 7c** and **Supplementary Fig. 9d-i**). In contrast, Na^+^ ions failed to traverse the pore despite having a smaller radius (1.2 Å), as expected for an anion selective pore. Consistent with the pore assuming an open state that is narrowest at the intracellular side, single channel recordings of GABA_A_Rs in membranes show reduced conductance upon mutation of the only polar side chain in the lowest two residue rings, γ2 2’ Ser, to non-polar alanine or valine^72^. Furthermore, the a1β3γ2_EM_ pore topology differs from the closed conformation of the related GlyR^6,7^, where the M2 helices were drawn together at the midpoint 9’ leucine residue ring to create a hydrophobic conductance barrier (2.8 Å diameter) (**Fig. 7b**). Nevertheless, the GABA_A_R open pore we observe is narrower than the pore of glycine-activated GlyR (8.8 Å diameter^6^) which, fittingly, possesses a higher conductance (60-90 pS^73^). From previous single channel studies wild type a1β2γ2 receptors in HEK cell membranes in the presence of saturating GABA concentrations occupy the open state 25 % of the time relative to 75 % for the desensitised state^74^. Presumably, the different experimental conditions imposed here, such as extraction from membranes into detergent micelles along with freezing in vitreous ice, have shifted the equilibrium in favour of the open state, resulting in its observation rather than the desensitised state.

**Figure 7.**
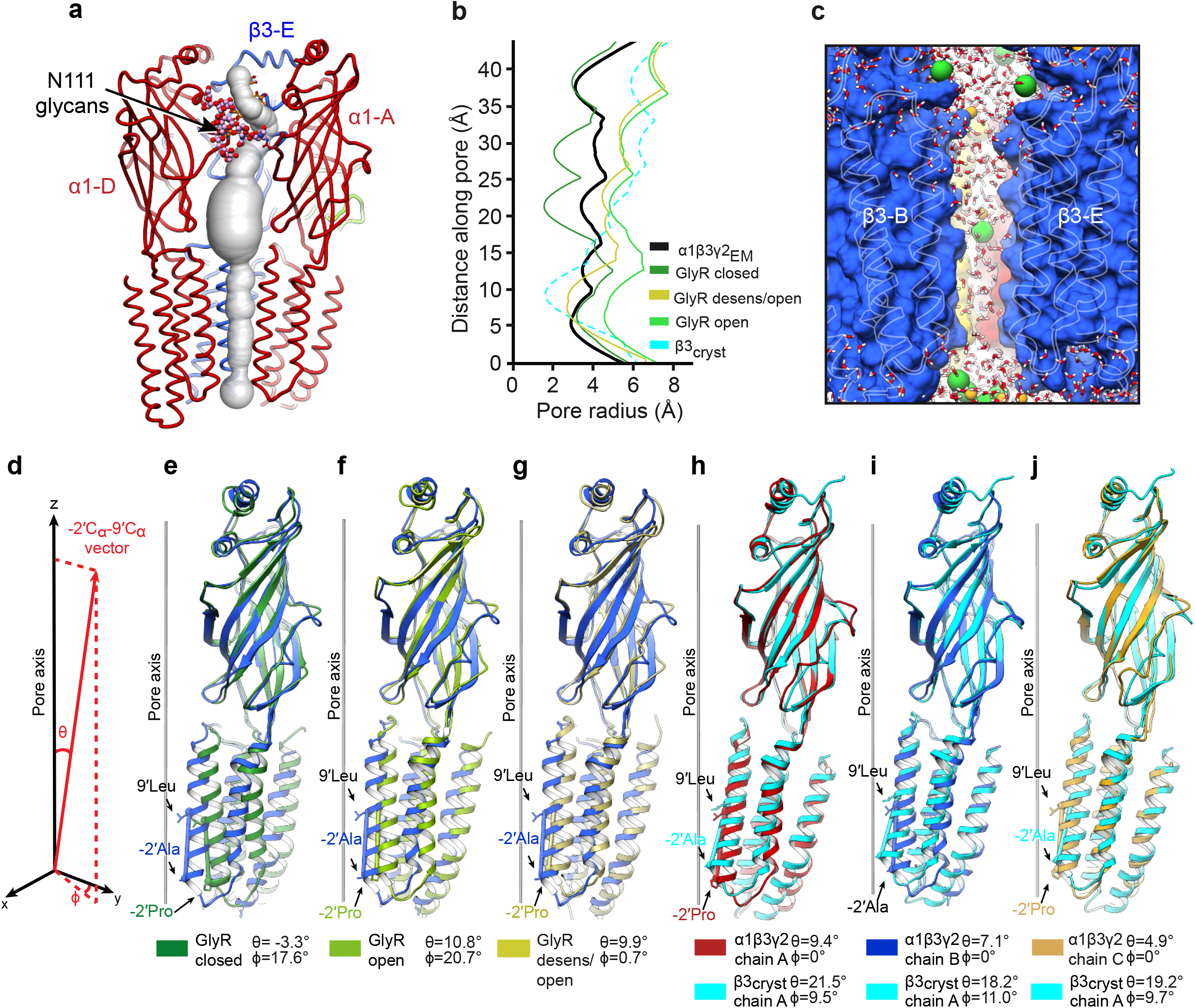
α1β3γ2_EM_ ion permeation pathway, open pore, and subunit conformations. (**a**) Solvent accessible tunnel through the vestibule between the α1 subunit Asn111-linked glycans and pore of α1β3γ2_EM_. β3-B and γ2-C subunits were hidden to visualise permeation pathway. The α1 Asn111-linked glycans are shown in ball and stick representation. (**b**) Plot of pore radii for α1β3γ2_EM_, desensitized GABA_A_R-β3_cryst_ (PDB ID: 4COF), glycine-bound GlyR (PDB ID: 5JAE), glycine/ivermectin-bound GlyR (PDB ID: 5JAF) and antagonist-bound GlyR (PDB ID: 5JAD). (**c**) Snapshot from a molecular dynamics simulation showing that the α1β3γ2_EM_ pore is fully hydrated and may be permeated by chloride ions. Pore viewed parallel to the membrane between two β3 subunits (represented as surface, white outlines are TMD helices). Chloride (green) and sodium (orange) ions are shown as van der Waals spheres whereas water molecules (white-red) are represented in bonds-only format. (**d**) Plot showing how the tilt (θ) and rotation (φ) angles are determined relative to the channel pore (z axis). M2 helices are defined as vectors between −2’ C_α_ and 9’ C_α_. Rotation angles for α1β3γ2_EM_ α1, β3 and γ2 subunits are set to zero. (**e-g**) Superposition between single subunits from GlyR in antagonist-bound, glycine-bound and glycine/ivermectin-bound conformations, respectively, and the α1β3γ2_EM_ β3 subunit. The ECD regions were used for structural alignment. A vector line between the M2 −2’ C_α_ and 9’C_α_ is drawn for each subunit. Tilt and rotation angles are shown for each pair. (**h-j**) Superposition between GABA_A_R-β3_cryst_ and α1β3γ2_EM_ α1, β3 and γ2 subunits, respectively, for TMD tilt and rotation analyses.

pLGICs in which proposed closed and open conformations of the same receptor have been determined, nAChR^75^ and GlyR^6^, reveal that pore opening arises from a relative ‘rocking’ motion between the ECD and the TMD regions. For α1β3γ2_EM_, the activated ECD base loops, in particular the β 1-2 loop and β6-7 loop (Cys-loop), are displaced away from the pore axis similarly to those of agonist-bound GlyR versus closed GlyR (**Supplementary Fig. 8n, o**). These base loops form contacts along the M2-M3 loop to concomitantly withdraw the TMD. Superposition of single ECDs from antagonist (strychnine), agonist (glycine) and glycine/ivermectin bound GlyRs on the α1β3γ2_EM_ β3 subunit reveal the impact of this ‘rocking’. Relative to the pore axis, closed GlyR M2 is over-straightened by θ = −3.3° and rotated clockwise by φ = 17.6°, which contrasts with the θ = 7.1° recline of the α1β3γ2_EM_ β3 subunit (**Fig. 7e**). As expected for the agonist-bound GlyR M2s, these assume similar reclines to α1β3γ2_EM_ β3, θ = 10.8° and 9.9°, respectively, with glycine alone bound GlyR also rotated anti-clockwise by φ = 20.7° (**Fig. 7f, g**).

To evaluate the possible transition motions from the α1β3γ2_EM_ open pore to the desensitized state^5,67^, we superposed an ECD from the desensitized GABA_A_R-β3_cryst_^5^ onto either α1, β3 or γ2 ECDs from the open pore α1β3γ2_EM_. This analysis revealed that the desensitized conformation also results from a combined tilting and rotation of subunit TMDs relative to the ECDs: the lower portions of the TMDs tilt 10-15° and rotate 9-11° relative to the α1β3γ2_EM_ subunits, moving towards the pore axis and closing the channel at the −2’ ring (**Fig. 7h-j**). A similar motion is observed for GlyR, in which the glycine plus ivermectin activated conformation undertakes an inward motion of the lower portions of the TMDs relative to the glycine bound alone activated state^6^.

## DISCUSSION

The α5_HOM_ and α5γ2_HET_ crystal structures reported here reveal the binding modes of the neutral ligand and BZD antagonist, flumazenil, and the partial PAM, bretazenil. These BZD sites retain 100 % residue identity to the wild type receptor site. Thus, these engineered templates serve as potent tools for future studies to unambiguously determine the binding modes of many of the small molecules previously found to bind the BZD site^24^. Of note, the binding affinities are in the range of one or two orders of magnitude lower for the BZDHET and BZDHOM sites, respectively. Nevertheless, the binding modes for flumazenil and bretazenil are maintained regardless of the site and are consistent with the previously established functional data discussed throughout the results. Importantly, introduction of a His to Arg substitution into the BZDHET site reproduced the same differential impact on flunitrazepam versus flumazenil binding as previously described for wild-type GABAA receptors^32,44^. Furthermore, superposition of these sites over the BZD site in α1β3γ2_EM_ reveal highly similar positioning of the secondary structure elements and relative appositions of the inter-subunit (P) and (C) faces. Thus, the lower affinities are not due to the sites being distorted or belonging to artefactual structures. Instead, differences likely reflect differing receptor energetics to assume favourable bound conformations for α5_HOM_ and α5γ2_HET_ versus wild type receptors in which two of the α-subunits are replaced by β-subunits.

Here we report activated homomeric and heteromeric GABA_A_R states bound by a BZD antagonist, and BZD and nanobody PAMs, respectively. These structures reveal fundamental insights into the activation and allosteric modulation processes, and lay the foundation to understand important aspects of GABA_A_R inhibitory neurotransmission. Furthermore, although BZDs are widely used to treat epilepsy, insomnia, anxiety, panic disorder and muscle spasm, they lack subtype selectivity and cause unwanted sedation, addiction, and motor and cognitive impairment^26^. Subtype selective drugs against the BZD site will ameliorate side effects and broaden the therapeutic repertoire to include treatments for autism, Down syndrome, neuropathic pain, schizophrenia and stroke^15–17,27,28^. We anticipate that the mechanistic insights and crystallographic platforms described here will expedite the arrival of structure-guided design of therapeutics to treat these disorders.

## ACKNOWLEDGMENTS

We thank staff at the Diamond Light Source beamline I04 for synchrotron assistance; K. Harlos and T. Walter for technical support with crystallization; R. Masiulyte for assistance in cryo-EM particle picking; G. Murshudov, I. Tickle and G. Bricogne for advice regarding anisotropic X-ray data processing; S. Scheres and D. Tegunov for advice regarding cryo-EM data processing; Y. Zhao for tissue culture advice; E. Beke for technical assistance during nanobody discovery; M. Duta for help at the Oxford Advanced Research Computing facility; D. Laverty for comments on the manuscript. This work was supported by the UK Biotechnology and Biological Sciences Research Council grants BB/M024709/1 (A.R.A. J.T.H. and P.S.M.) and BB/N000145/1 (S.J.T. and M.S.P.S.); UK Medical Research Council grants MR/L009609/1 (A.R.A.), MC_UP_1201/15 (A.R.A. and S.M.) and MC_U105185859 (M.M.B. and S.C.); Wellcome Trust studentships 084655/Z/08/Z (S.M.) and 105247/Z/14/Z (S.S.); Human Frontier Science Program grant RGP0065/2014 (A.R.A.) and long-term postdoctoral fellowship LT000021/2014-L (T.M.); Cancer Research UK grant C20724/A14414 (T.M.); European Research Council grant 649053 (J.T.H.); EPSRC grant EP/R004722/1 (M.S.P.S.) and Wellcome grant 208361/Z/17/Z (M.S.P.S.). We thank INSTRUCT, part of the European Strategy Forum on Research Infrastructures and the Research Foundation-Flanders (FWO) for funding nanobody discovery. Further support from the Wellcome Trust Core Award 090532/Z/09/Z is acknowledged. The OPIC electron microscopy facility was founded by a Wellcome Trust JIF award (060208/Z/00/Z) and is supported by a WT equipment grant (093305/Z/10/Z).

## AUTHOR CONTRIBUTIONS

Molecular biology, protein expression, purification and crystallization, radioligand binding assays, whole cell electrophysiology: PSM; X-ray data collection: A.R.A., P.S.M., T.M.; X-ray data processing: A.R.A., P.S.M.; cryo-EM data collection: S.M., A.K., Z.S., P.S.M., J.T.H.; cryo-EM data processing: S.M., J.T.H; cryo-EM model building: S.M., L.D.C., S.L., B.F., F.D.M.; small-molecule docking: T.M.; human genome bioinformatics: S.C., M.M.B.; molecular dynamics simulations: S.R., G.K., S.J.T., M.S.P.S; software and hardware: J.M.D, A.S., R.M.E.; nanobody generation: E.P., J.S.; SPR: S.M.; slice electrophysiology: S.H., T.G.S. The manuscript was written by P.S.M, S.M. T.M. and A.R.A, with input from all co-authors.

## METHODS

### Construct design, α5_HOM_ and α5γ2_HET_

Details of the α5 subunit constructs design, including protein sequences, are shown in **Supplementary Figure 1**. The chimeric γ2-ECD:α1-TMD subunit of α5γ2_HET_ comprises mature sequence (Uniprot P18507) γ2 residues 39 to 232 (QKSDD…DLSRR) appended to α1 (Uniprot P62813) from 223 to 455 (IGYFVI…PTPHQ) with a single β3 substitution, (α1 P280A). The α5 intracellular M3-M4 loop amino acids 316-392 (RGWA…NSIS) (Uniprot P31644) and the α1 intracellular M3-M4 loop amino acids 313-390 (RGYA…NSVS) were substituted by the SQPARAA sequence^5,40^ to enhance the recombinant protein yield and facilitate crystallisation. Constructs were cloned into the pHLsec vector^76^, between the N-terminal secretion signal sequence and either a double stop codon or a C-terminal 1D4 purification tag derived from bovine rhodopsin (TETSQVAPA) that is recognised by the Rho-1D4 monoclonal antibody (University of British Columbia)^77,78^.

### Construct design, α1β3γ2_EM_

The protein sequences used were: human GABA_A_R α1 (mature polypeptide numbering 1-416, QPSL…TPHQ; Uniprot P14867), human GABA_A_R β3 (mature polypeptide numbering 1-447, QSVN…YYVN; Uniprot P28472), human GABA_A_R γ2 (mature polypeptide numbering 1-427, QKSD…YLYL; Uniprot P18507). These constructs were cloned into the pHLsec vector^76^, after the N-terminal secretion signal sequence and before a double stop codon unless stated otherwise below. The a1 intracellular M3-M4 loop amino acids 313-391 (RGYA…NSVS) were substituted by the SQPARAA sequence^5,40^. Four β3 ECD residues were replaced by β2 subunit residues (Gly171Asp, Lys173Asn, Glu179Thr, Arg180Lys), which block homomer assembly^79^, and the β3 intracellular M3-M4 loop amino acids 308-423 (GRGP…TDVN) were substituted by a modified SQPARAA sequence containing the *E.coli* soluble cytochrome B562RIL^41^ (BRIL, amino acids 23-130, ADLE…QKYL, Uniprot P0ABE7) to give an M3-M4 loop of sequence SQPAGT-BRIL-TGRAA. The γ2 intracellular M3-M4 loop amino acids 323-400 (NRKP…IRIA) were substituted by the SQPARAA sequence, and a C-terminal GTGGT linker followed by a 1D4 purification tag derived from bovine rhodopsin (TETSQVAPA) that is recognised by the Rho-1D4 monoclonal antibody (University of British Columbia)^77,78^. Nb38 was identified from a previously described nanobody library^20^.

### Large-scale expression and purification of a5_HOM_, a5γ2_HET_ and a1β3γ2_EM_

Twenty-litre batches of HEK293S-GnTI^−^ cells (which yield proteins with truncated N-linked glycans, Man_5_GlcNAc_2_^80,81^) were grown in suspension to densities of 2 × 10^6^ cells ml^−1^ in Protein Expression Media (PEM, Invitrogen) supplemented with L-glutamine, non-essential amino-acids (Gibco) and 1% foetal calf serum (Sigma-Aldrich). Typical culture volumes were 200 ml, in 600 ml recycled media bottles, with lids loose, shaking at 130 rpm, 37°C, 8 % CO_2_. For transient transfection, cells from 1 litre of culture were collected by centrifugation (200 *g* for 5 min) and resuspended in 150 ml Freestyle medium (Invitrogen) containing 3 mg PEI Max (Polysciences) and 1 mg plasmid DNA, followed by a 4 h shaker-incubation in a 2 litre conical flask at 160 rpm. For α5γ2_HET_ DNA plasmids were transfected at 9:1 ratio (i.e. 0.9:0.1 mg) α5 construct DNA without a 1D4 tag to the chimera γ2-ECD:α1-TMD with a 1D4 purification tag. For α1β3γ2_EM_ DNA plasmid ratios were 1:1:0.5, respectively. Subsequently, culture media were topped up to 1 litre with PEM containing 1 mM valproic acid and returned to empty bottles. Typically, 40-70 % transfection efficiencies were achieved, as assessed by control transfections with a monoVenus-expressing plasmid^82,83^. 72 h post-transfection cell pellets were collected, snap-frozen in liquid N2 and stored at −80 °C.

Cell pellets (approx. 200g) were solubilised in 600 ml buffer containing 20 mM HEPES pH 7.2, 300 mM NaCl, 1 % (v/v) mammalian protease inhibitor cocktail (Sigma-Aldrich, cat. P8340) and 1.5 % (w/v) dodecyl 1-thio-β-maltoside (DDTM, Anatrace) for α5_HOM_ and α1β3γ2_EM_ or 1.5 % (w/v) decyl β-maltoside (DM, Anatrace) for α5γ2_HET_, for 2 hours at 4 °C. Insoluble material was removed by centrifugation (10,000 g, 15 min). The supernatant was diluted 2-fold in a buffer containing 20 mM HEPES pH 7.2, 300 mM NaCl and incubated for 2 hr at 4 °C with 10 ml CNBr-activated sepharose beads (GE Healthcare) pre-coated with 50 mg Rho-1D4 antibody (3.3 g dry powdered beads expand during antibody coupling to approximately 10 ml). Affinity-bound samples were washed slowly by gravity flow over 2 hours at 4 °C with 200 ml buffer containing 20 mM HEPES pH 7.2, 300 mM NaCl, and either 0.1 % (w/v) DDTM (approximately 20 x CMC) for α5_HOM_, or 0.2 % (w/v) DM (approximately 3 x CMC) α5γ2_HET_, or for α1β3γ2_EM_ 0.1 % (w/v) DDTM and 0.01 % (w/v) porcine brain polar lipid extract (141101C Avanti; chloroform was evaporated under argon then 100 mg lipid film was dissolved in 10 ml 10 % (w/v) DDTM (1000 mg) in water and stored at −80 C until needed). Beads were then washed in a second round of buffer: 20 mM HEPES pH 7.2, 300 mM NaCl, and either 0.01 % (w/v) DDTM (approximately 3 x CMC) for α5_HOM_ or 0.2 % (w/v) DM (approximately 3 x CMC) for α5γ2_HET_, or for α1β3γ2_EM_, 1mM GABA, 0.01 % (w/v) DDTM (approximately 4 x CMC), 0.001 % (w/v) porcine brain polar lipid extract. Protein samples were eluted overnight in 15 ml buffer containing 15 mM HEPES pH 7.2, 225 mM NaCl, 500 μM TETSQVAPA peptide (Genscript), and corresponding detergents. The eluate was centrifuged (30,000g, 15 min) and the supernatant was concentrated by ultrafiltration to 1-2 ml at 1-5 mg/ml using 100-kDa cut-off membranes (Millipore). The concentrated sample was centrifuged (30,000 g, 15 min) and the supernatant was aliquoted in 0.5-1.5 mg protein per 0.7 ml aliquots and either snap-frozen for storage at −80 °C or gel filtrated as appropriate. A single aliquot was loaded onto a Superose 6 10/300 Increase gel filtration column (GE Healthcare) equilibrated in 10 mM HEPES pH 7.2, 150 mM NaCl, and either: 0.007 % (w/v) DDTM, 50 μM flumazenil, 50 μM pregnanolone for a5_HOM_ or 0.2 % (w/v) DM, 50 μM bretazenil for a5γ2_HET_, or for a1β3γ2_EM_, 1 mM GABA, 0.007 % (w/v) DDTM. The peak fractions were approximately 0.5 mg/ml. The fractionated protein was concentrated by ultrafiltration to 3-5 mg/ml, using 100 kDa cut-off membranes (Millipore), for crystallisation trials. Typical final yields were 0.1-0.2 mg protein per litre of cells grown in suspension (10 g cell pellet). In the case of a1β3γ2_EM_, Nb38 was added to a1β3γ2 at 4-fold molar excess and the complex was concentrated by ultrafiltration to 2.5 mg/ml, using 100 kDa cut-off membranes (Millipore).

### Nb38 purification

Nb38 was produced and purified in milligram quantities from WK6su^−^ *E. coli* bacteria^84^. Bacteria were transformed with ∼200 ng of nanobody expression plasmid pMESy4 containing the nanobody of interest and selected on Lysogeny broth (LB)-agar plates containing 2% glucose and 100 μg/ml ampicillin. 2-3 colonies were used for preparing a preculture, which was used to inoculate 0.5 L Terrific broth (TB) cultures supplemented with 0.1 % glucose, 2 mM MgCl_2_ and 100 μg/mL ampicillin. Cultures were grown at 37 °C until their OD_600_ reached 0.7, at which point Nb38 expression was induced with 1 mM IPTG. After induction, cells were grown at 28 °C overnight and harvested by centrifugation (20 min, 5000g). Nanobodies were released from the bacterial periplasm by incubating cell pellets with an osmotic shock buffer containing 0.2 M Tris pH 8.0, 0.5 mM EDTA, and 0.5 M sucrose. The C-terminally His6-tagged Nb38 was purified using nickel affinity chromatography (binding buffer: 50 mM HEPES pH 7.6, 1 M NaCl, 10 mM imidazole; elution buffer: 50 mM HEPES pH 7.6, 0.2 M NaCl, 0.5 M imidazole), followed by size-exclusion chromatography on a Superdex 75 16/600 column (GE Healthcare) in 10 mM Hepes pH 7.6, 150 mM NaCl. Nb38 stocks were concentrated to 5-10 mg/mL, snap-frozen in liquid nitrogen and stored at −80 °C.

### Crystallization and data collection

α5γ2_HET_ and α5_HOM_ contain 15 N-linked glycosylation sites each, bringing a considerable extra volume, flexibility and potential occupancy heterogeneity. Therefore, prior to crystallization, concentrated protein samples (6 mg/ml α5γ2_HET_ and 4 mg/ml α5_HOM_) were incubated with 0.01 mg ml^−1^ endoglycosidase F1^85^ for 2h at room temperature. Sitting drop vapour diffusion crystallization trials were performed in 96-well Swisssci 3 well crystallisation plates (Hampton Research), at three ratios: 200 nl protein plus 100 nl reservoir, 100 nl protein plus 100 nl reservoir, 100 nl protein plus 200 nl reservoir. Drops were dispensed by a Cartesian Technologies robot^86^, and plates were maintained at 6.5 °C in a Formulatrix storage and imaging system. In the case of α5γ2_HET_, crystals appeared in a range of conditions^87^ within 1-28 days, with the best diffracting crystals (to ∼2.5 Å resolution) taking 4 weeks to grow in: 22 % poly-ethylene (PEG) 400, 0.37 M potassium nitrate, 0.1 molar 2-(N-morpholino)ethanesulfonic acid (MES) pH 6.5. For α5_HOM_, crystals also grew in a range of conditions, typically within 2 weeks, and in the first instance diffracted up to intermediate resolution (>5 Å). Following additive-based optimization (MemAdvantage, Molecular Dimensions), crystals diffracting to ∼2.6 Å resolution were identified, grown in: 19 % PEG 1000, 0.1 M sodium chloride, 0.15 M ammonium sulphate, 0.1 M MES pH 6.5, 2.5 mM sucrose monodecanoate (sucrose monocaprate). Crystals were cryoprotected by soaking in reservoir solution supplemented with 30 % ethylene glycol, and then cryocooled in liquid nitrogen. Diffraction images were collected at the Diamond Light Source beamline I04, λ=0.9795 Å, 0.1° oscillation (bretazenil-bound α5γ2_HET_) and 0.2° oscillation (flumazenil-bound α5_HOM_), on a Pilatus 6M-F detector. X-ray data were indexed, integrated and scaled using the HKL2000 package^88^. Diffraction from both α5γ2_HET_ and α5_HOM_ crystals was severely anisotropic, therefore scaled but unmerged data were processed with STARANISO^89^, allowing for the anisotropic diffraction cut-offs to be applied before merging with Aimless^90,91^, within the autoPROC toolbox^92^. Upon ellipsoidal truncation, resolution limits were 2.33 Å, 3.15 Å and 3.73 Å (in the −0.022 *a*\* + *c*\*, *b*\* and 0.945 *a*\* - 0.327 *c*\* directions, respectively) for α5γ2_HET_, and 2.49 Å, 3.13 Å and 4.63 Å (in the 0.872 *a*\* - 0.490 *c*\*, *b*\* and 0.842 *a*\* + 0.540 *c*\* directions, respectively) for α5_HOM_. Data collection and merging statistics are detailed in the **Table 1**.

### Structure determination, refinement and analysis

α5γ2_HET_ and α5_HOM_ structures were solved by molecular replacement using the human GABA_A_R-β3_crys_t homopentamer^5^ (PDB ID: 4COF) as a search model in Phaser^93^. Polypeptide chains were traced using iterative rounds of manual model building in Coot^94^ and refinement in BUSTER-TNT^95^, Refmac^96^ and Phenix^97,98^. Automated X-ray and atomic displacement parameter (ADP) weight optimisation, and torsion angle non-crystallographic symmetry (NCS) restraints, were applied. Ligand coordinates and geometry restraints were generated using the grade server^99^. The α5γ2_HET_ and α5_HOM_ models contain one homopentamer per asymmetric unit. Crystal packing impaired map quality in regions where ECD from certain subunits were near of detergent micelles of neighbouring molecules. Nevertheless, complete polypeptide chains could be built, with the exception of 14 N-terminal α5 residues (QMPTSSVKDETNDN), 22 N-terminal γ2 residues (QKSDDDYEDYTSNKTWVLTPKV) and the C-terminal purification tags, presumably disordered. Strong additional electron density peaks were clearly visible in the BZD_HOM_ and BZD sites, that could be unambiguously assigned to flumazenil in a5_HOM_ and bretazenil in a5γ2_HET_, respectively, based on shape, coordination and refinement statistics. Furthermore, electron density corresponding to five pregnanolone molecules, one per inter-subunit interface, could be observed at the TMD interfaces of a5_HOM_, as previously described^20^, and five well-ordered detergent molecules (decyl β-maltoside) could be built inside the a5γ2_HET_ pore region. The a5 and γ2 extracellular regions have three N-linked glycosylation sites each, and we could observe clear electron density for six NAG moieties in a5_HOM_ and five in a5γ2_HET_, the others being disordered. Stereochemical properties of the models were assessed in Coot^94^ and Molprobity^100^. Refinement statistics are provided in the **Table 1**. Protein geometry analysis revealed no Ramachandran outliers, 97.96% residues in favoured regions and 2.04% residues in allowed regions for a5_HOM_, and one Ramachandran outlier (0.06%), 95.97% residues in favoured regions and 4.03% residues in allowed regions for a5γ2_HET_. Molprobity clash scores after adding hydrogen atoms is 9.16 (99^th^ percentile) for a5_HOM_ and 11.06 (94^th^ percentile) for a5γ2_HET_. Overall Molprobity scores are 1.50 (100^th^ percentile) for a5_HOM_ and 1.84 (98^th^ percentile) for a5γ2_HET_. Structural alignments were performed in PyMOL using the align function. Structural figures were prepared with the PyMOL Molecular Graphics System, Version 1.8, Schrödinger, LLC., and the UCSF Chimera package, developed by the Resource for Biocomputing, Visualization, and Informatics at the University of California, San Francisco^101^.

### Electron microscopy and image processing

CryoEM samples were prepared using C-flat^™^ Holey Carbon grids (R2/1, 200 mesh, 53% of collected images) and UltraAuFoilTM grids (R1.2/1.3, 200 mesh, 47% of collected images). Carbon substrate grids were glow discharged for 10 s, then 3.5 μL of protein sample (4.2 mg/mL) was applied for 15 s. The sample was blotted for 3 s using VitroBot Mark IV (FEI) and flash-frozen in liquid ethane. Gold substrate grids were glow discharged for 15 s and 2.5 μL sample (2.5 mg/mL) was applied for 30s, which was followed by 7.5 s blotting. In both cases, blotting was performed at room temperature and 90-100% humidity. All cryo-EM data were collected using a Tecnai F30 Polara electron microscope (FEI) operating at 300 kV, fitted with a K2 Summit direct electron detector (Gatan) and a GIF-Quantum energy filter (Gatan). SerialEM was used to manually record zero energy-loss (20 eV slit) images at a calibrated magnification of 37,000x (1.35Å/pix) in counting mode. Images were recorded as movies consisting of 47 frames with total dose of 38 e^−^/ Å^2^ and exposure of 14.1s. Nominal defocus values ranged from −2.0 to −3.5 μm.

Classification of α1β3γ2_EM_ particles in the decylmaltoside neopentylglycol, (DMNG) detergent revealed that the particles had a ‘collapsed’ TMD region, with the γ2 TMD occupying the central pore (**Supplementary Fig. 4a**). We discarded this sample, considering it to be a sample preparation artefact, presumably caused by extraction of the receptor from its native cell bilayer into DMNG alone. Preparation of a second sample in a different detergent, dodecyl 1-thio-β-maltoside (DDTM) and in the presence of porcine brain polar lipid extract (see methods section on Large-scale expression and purification) yielded the predominant particle population with TMDs pseudo-symmetrically arranged around the pore, i.e. the γ2 TMD was no longer ‘collapsed’ into central pore. This sample was used for all subsequent processing. A total of 8,548 movies were motion-corrected at micrograph level with UCSF MotionCorr^102^ and CTFFIND4^103^ was used to estimate the contrast function parameters. Data processing was performed using RELION software package^104^. First, ∼5,000 particles were picked using EMAN2^105^, grouped in RELION using reference-free 2D classification and class averages were used as references for RELION automated particle picking in the same software. False-positives were manually removed leading to a dataset of 525,124 particles. Reference-free 2D classification was performed and particles belonging to classes showing GABA_A_R features were selected (436,320 particles). An initial 3D model was generated from the GABA_A_R-β3_cryst_ structure^5^ (PDB ID 4COF) and low-pass filtered to 40 Å resolution. Particles were first oriented in 3D by imposing C5 symmetry, and further classified in 3D imposing no symmetry (C1) by only allowing rotation around the symmetry axis. The best class showing features of two BRILs and two nanobodies was selected and used as a new initial reference model. Particles selected from 2D classes were 3D-classified into 10 classes using C1 symmetry. Particles assigned to the best 3D classes (186,786 particles) were used for particle ‘polishing’ step in RELION, where particle motion and radiation damage for particles from each movie frame was estimated. ‘Gold standard’ refinement of the resulting ‘shiny’ particles resulted in a 5.65 Å map. Movie motion correction with MotionCor2^106^ using 25 patches and dose-weighting scheme improved the resolution of the refined map to 5.25 Å. These particles were further 3D-classified into 10 volumes. Particles assigned to the best 9 classes (165,621 particles) were combined to yield a map of 5.17 Å. The resolution was estimated using *relion post-process* with the FSC criteria of 0.143. The final unsharpened and unfiltered map was globally autosharpened using *phenix.auto_sharpen^107^* (5.17 Å high-resolution cut-off) to maximise the map detail while maintaining the connectivity of the map.

### Model building and refinement of α1β3γ2_EM_

Crystal structures (2.9 Å β3_cryst_ (PDB ID 4COF), 2.5 Å a5γ2HET (PDB ID XXXX) and 3.2 Å a5TMD^20^ (PDB ID 5O8F)) were used to build the GABA_A_R α1β3γ2_EM_ heteromer and Nb38 models into the 5.17 Å map. First, the a5 subunit and Nb25^20^ (5O8F) structures were manually mutated to match the a1 and Nb38 sequences, respectively. TMD of the γ2 subunit was prepared by mutating a5 TMD (5O8F) residues to match the γ2 sequence. The Nb38 model and the corresponding ECDs and TMDs of α1, β3 and γ2 subunits were docked into the cryo-EM map using Chimera. Map was cut around each subunit and nanobody (4 Å radius) using Chimera. Rosetta-CM^108^ was used to refine the models into the resulting cryo-EM densities to improve model geometry and fitting in the density. N-linked glycan models from α5_TMD_ crystal structure (5O8F) were docked into the cryo-EM map and added to the model. Rotamers were manually adjusted to match the high resolution structures; where no prior information was available, the most common rotamers were chosen. The model was further optimised by rounds of manual correction in Coot^94^ and iterative refinement in real space with *phenix_real_space_refine^109^* using secondary structure and NCS restraints. The final model contains α1 subunit residues 12-312 (TTVF…YFTK) and 391-414 (KIDR…YWAT); β3 subunit residues 10-307 (SFVK…YIFF) and 422-447 (AIDR…LYYV); γ2 subunit residues 27-322 (VTVI…YFVS) and 401-424 (KMDS…YWVS) and Nb38 residues 1-123 (QVQL…TVSS). For model validation, the final model coordinates were randomly displaced by 0.2 Å and then this model was refined with *phenix_real_space_refine* against one of the half-maps produced by RELION^104^. FSC curves were then calculated between the refined model and half-map used for refinement (‘work’) and between the second half-map, not used for refinement, (‘free’). No significant separation between FSC_wor_k and FSC_free_ curves was observed, indicating the model was not over-refined. The stereochemistry of the final model was evaluated using MolProbity^110^. Figures were prepared using UCSF Chimera package, developed by the Resource for Biocomputing, Visualization, and Informatics at the University of California, San Francisco^101^. Structural alignments were performed in SHP^111^. Pore dimensions were analysed using the Coot implementation of Hole^112^ and with CAVER^113^ PyMOL plugin with a probe radius of 2.5 Å.

### Small molecule docking

Molecular docking of small molecules to a5γ2_HET_ and a5_HOM_ crystal structures was performed using AutoDock Vina^114^. Stereochemistry of small molecules was optimised using Grade webserver^115^. Structures of the receptor were kept rigid during docking. The region selected for docking encompassed whole benzamidine-binding pocket.

### Molecular dynamics simulations

The protein portion of the receptor structure, either full length (for uniform electric field simulations) or with only the transmembrane domain included (for water equilibrium simulations), was embedded within a POPC lipid bilayer, in a simulation box containing an aqueous solution of Cl^−^ and Na^+^ ions at 0.5 M or 0.15 M concentration, respectively, following a previously established protocol^116^ and employing the TIP4P water model^117^. Simulations were performed using GROMACS version 5.1^118,119^, and the OPLS united-atom force field^120^. The temperature and pressure were maintained at 37 °C and 1 bar, respectively, using the velocity-rescale thermostat^121^ in combination with a semi-isotropic Parrinello and Rahman barostat^122^, with coupling constants of τ_T_ = 0.1 ps and τ_P_ = 1 ps. A Verlet cut-off scheme was applied, and long-range electrostatic interactions were measured using the Particle Mesh Ewald method^123^. Bonds were constrained using the LINCS algorithm^124^, and an additional harmonic restraint at a force constant of 1000 kJ mol^−1^ nm^−2^ was placed on protein backbone atoms to preserve the conformational state of the original, experimentally determined structure. The integration time-step was 2 fs. For the water distribution and free energy profiles: 50 ns repeats were each initiated from an independently assembled simulation system. Free energy profiles for water permeation along the channel pore were derived (using the CHAP channel annotation package, http://www.channotation.org/) from the equilibrium distribution of water molecules upon simulation through an inverse Boltzmann calculation-based method^70^. Pore radius profiles (i.e. radii at positions along the pore axis) were estimated using the programme HOLE^112^. To determine the behaviour of ions a +0.12 V potential difference was imposed across the membrane during 100 ns simulations by setting a linear potential difference of 0.5 V across the z-length of the simulation box. The conductance was estimated based on the number of Cl^−^ ions passing through the channel.

### Radioligand binding experiments

GABA_A_R constructs containing a single BZD site (α1β3γ2_EM_, α5γ2_HET_Δ and α5β3γ2_WT_) at 2 nM and five sites (α5γ2_HET_ and α5_HOM_) at 0.4 nM, were used, in 10 mM HEPES pH 7.2, 150 mM NaCl, and: 0.05 % (w/v) DDTM, 0.005 % (w/v) porcine brain polar lipid extract; 0.2 % (w/v) DM detergent for α5γ2_HET_; 0.05 % (w/v) detergent (decylmaltoside neopentylglycol (DMNG) 5:1 (molar ratio) cholesterol hemisuccinate (CHS) for α5β3γ2_WT_; 0.05 % n-dodecyl-beta-maltoside (DDM) for α5γ2_HETΔ_ and α5_HOM_. Samples were incubated with WGA YSI beads (bind N-linked glycans, beads at 2 mg/ml, Perkin Elmer) for 30 minutes at 4 °C under slow rotation. 50 μl aliquots of the GABA_A_ receptor-bead mix were added to 50 μl aliquots of 2 x radioligand ([^3^H]-flunitrazepam or [^3^H]-flumazenil) concentrations ranging from 0.06-2000 nM (Perkin Elmer) in Serocluster 96-Well ‘U’ Bottom plates (Corning) and incubated for 60 minutes at room temperature (20-22 °C) and [^3^H] cpm were determined by scintillation proximity assay using a Microbeta TriLUX 1450 LSC. The same ligand binding assay was performed in the presence of 50 μM flumazenil to ascertain the non-specific binding (NSB), which was subtracted from the total radioligand cpm to obtain the specific binding values. [^3^H]-flunitrazepam binding affinity (K_d_) was calculated in OriginPro2015 using the one-site binding curve fit equation (*y* = *B_max_*\**x*/(*k1*+*x*)), or two-site binding curve fit equation (*y* = *B_max1_*\**x*/(*k1*+*x*) + *B_max_2*\**x*/(*k2*+*x*)), or using the Hill equation (*y* = *B_max_*\**x^n^*/(*k1^n^*+*x^n^*) where *B_max_* values are maximal binding for each site and *k1* and *k2* are *K_d_* for each site, *n* is Hill slope, x is ligand concentration, *y* is proportion of binding. Displacement curves were performed by adding ligand (bretazenil or diazepam) over the concentration range 1-50000 nM to aliquots of GABA_A_ receptor-bead mix for 30 minutes, then adding this to aliquots of radioligand ([^3^H]-flumazenil or [^3^H]-flunitrazepam, respectively) at final concentrations corresponding to approximately 10 x *K_d_*. Diazepam displacement curves were plotted on log concentration axis and fitted using the logistic equation (*y* = *A2*+(*A2*-*A1*)/*1*+(*x*/*x0*)^*p*) where *A2* and *A1* are maximal and minimal binding respectively, *x0* is IC_50_ and *p* is the Hill coefficient. *IC_50_* values of displacement curves were converted to *Ki* values according to the Cheng-Prusoff equation^125^, *Ki* = *IC_50_/1*+(*[L]/K_d_*) referring to the [^3^H]-flumazenil *K_d_* and the bretazenil *IC_50_*, and where *L* is the concentration of [^3^H]-flumazenil used in the displacement assay.

### Thermostability binding experiments

Information regarding the thermostability of a detergent-solubilised protein can be determined by heating protein samples over a range of temperatures for equal time periods and then measuring the reduction in the intensity of the monodisperse SEC profile for each protein sample^126^. With increasing temperature an increased proportion of protein is denatured, aggregates and is lost from the monodisperse peak when the protein is subsequently run on SEC. A measure of protein stability can then be obtained by plotting the decay in peak UV absorbance against increasing temperature, for example to obtain a 50 % melting temperature (T_m_). Purified a1β3γ2_EM_ at 0.02 mg ml^−1^ (100 nM) in 150 mM NaCl, 10 mM HEPES pH 7.2, 0.007 % DDTM (w/v) was separated into 50 μl aliquots in PCR tubes, and heated at a range of temperatures from 30-80 °C for 1 hour. Samples were then run on a high-performance liquid chromatography system with automated microvolume loader (Shimadzu) through a Superdex 200 Increase 3.2/300 column (GE Healthcare) maintained in 300 mM NaCl, 10 mM HEPES pH 7.2, 0.007 % DDTM (w/v). Monodisperse peak reduction with increasing temperature was measured relative to an unheated control sample maintained at 4 °C.

Importantly, because some drugs when bound thermostabilise detergent-solubilised protein^126^, the thermostability assay offers an efficient strategy to measure protein sensitivity to drugs in the detergent-solubilised environment. Purified GABA_A_R Cryo-EM construct was separated into PCR tubes, supplemented with picrotoxin at a range of concentrations and heated at Tm30 % (the temperature at which the monodisperse peak was reduced by 70 %) for 1 hour. Afterwards samples were run on a high-performance liquid chromatography system with automated micro-volume loader (Shimadzu) through a Superdex 200 3.2/300 column (GE Healthcare) maintained in 300 mM NaCl, 10 mM HEPES pH 7.2, 0.007 % DDTM. Drug dose-response curves were generated by plotting UV absorbance against drug concentration.

Surface plasmon resonance analysis (SPR). SPR measurements were performed using a Biacore T200 (GE Healthcare) machine at 25 °C. All reagents and consumables for SPR were purchased from GE Healthcare. The carboxyl groups on CM5 chip flow channels were activated with a 10-minute injection of a 1:1 mixture of 0.1 M N-hydroxysuccinimide (NHS) and 0.4 M 1-ethyl-3-dimethylaminopropyl-carbodiimide (EDC). Streptavidin (Sigma-Aldrich) was covalently bound by a 5-minute injection until an immobilization level of 5000 RU was reached. Free activated carboxyl groups were quenched with a 10-minute injection of 1 M ethanolamine-HCl pH 8.5. The working flow-cell was functionalized by injecting GABA_A_ receptor electron cryo-microscopy construct containing a biotinylated C-terminus containing a biotin ligase recognition sequence, until 350-375 RU were reached, whereas the reference cell containing streptavidin was left unmodified. Running buffer contained 10 mM HEPES pH 7.5, 150 mM NaCl, 0.007% DMNG:CHS (5:1, w/w). For experiments testing GABA effect on Nb38 binding, 1 mM GABA was supplemented to the running buffer. Measurements were performed by injecting nanobodies at concentrations ranging from 2.5 to 40 nM (two-fold serial dilutions) during a single cycle. For reliable single cycle kinetics (SCK) data fitting, the final dissociation phase was set to 15 minutes. Biacore T200 evaluation software was used to analyse all the SCK data. A 1:1 binding model was used to fit the experimental results.

### HEK cell preparation and electrophysiology

One day prior to experiments, 8 ml of Dulbecco’s Modified Eagle Medium (DMEM) was pre-incubated for 10 min at room temperature with 96 μl lipofectamine2000 (Thermofisher) and 48 μg plasmid DNA, then added to a single T175 cm^2^ flask containing HEK293T cells (30-50 % confluency) and 2 ml DMEM (supplemented with 10 % fetal calf serum, L-Gln and non-essential amino acids). After 3 hrs this media was removed and replaced by DMEM supplemented with 10 % fetal calf serum. For GABA_A_R heteromers, pHLsec plasmids containing human cDNA constructs were mixed in 1:1:2 ratio (a:β:γ), supplemented with 3% plasmid encoding enhanced green fluorescent protein (EGFP) to assess transfection efficiency. The electron cryo-microscopy a1β3γ2 construct used was as described in ‘Construct design’. The wild-type cDNA inserts used for heteromeric receptor expression were as follows: human GABA_A_R a1 mature protein sequence (a1 Uniprot P14867 entry, Gln28 is Gln1, 1-429 QPSL…PTPHQ) and human β3 mature protein sequence (β3 Uniprot P28472 entry isoform 1, Gln26 is Gln1, 1-448 QVSN…LYYVN) cloned into the pHLsec vector^34^ between the N-terminal secretion signal sequence and a double stop codon; Human GABA_A_R γ2 mature protein sequence (γ2 Uniprot P18507 isoform 1 entry Gln40 is Gln1 1-428 QKSDD…SYLYL) cloned into the pHLsec vector^34^ between the N-terminal secretion signal sequence followed by streptavidin binding protein (MDEKTTGWRGGHVVEGLAGELEQLRARLEHHPQGQREP) and a C-terminal 1D4 purification tag. Transfection efficiencies were typically 50-80 % (cells expressing EGFP, as estimated by fluorescence microscopy). Eighteen to twenty-four hours later cells were washed with phosphate buffered saline, incubated in 4 ml TrypLE (Gibco) for 7 min at 37 °C, suspended in 21 ml DMEM supplemented with 10 % fetal calf serum and L-Gln, centrifuged at 100 *g* for 1.5 min, then suspended in 50 ml Freestyle 293 Expression Medium (Gibco) and placed in a shaking incubator (130 rpm, 37°C, 8 % CO_2_) for 30 min. 25 ml cell suspension was then centrifuged at 100 *g* for 1.5 min, and suspended in 4 ml external recording solution. This solution contained (mM): 137 NaCl, 4 KCl, 1 MgCl_2_, 1.8 CaCl_2_, 10 HEPES, and 10 D-Glucose, pH 7.4 (≈ 305 mOsm). The internal recording solution contained (mM): 140 CsCl, 5 NaCl, 1 MgCl_2_, 10 HEPES, 5 EGTA, 0.2 ATP, pH 7.35 (≈ 295 mOsm). Electrophysiological recordings were performed at room temperature using an Ionflux16 (Molecular Devices) in ensemble mode, with series resistance compensation set at 80 % and cells held at −60 mV. Diazepam and picrotoxin (Sigma-Aldrich) were dissolved in DMSO as 100 mM and 1 M stocks respectively prior to dilution in external recording solution. Diazepam and Nb38, or picrotoxin, were coapplied with EC_10-15_ GABA or EC_70_ GABA doses respectively to generate dose response curves. Expression of heteromeric receptors as assemblies of αβγ subunits was confirmed by response to GABA, which requires co-assembly of α1 and β3 subunits, and efficient inclusion of the γ2 subunit into αβγ heteromers was verified by measuring low sensitivity to 100 μM Zn^2+^ inhibition, defined as less than 50 % inhibition of an EC_50_ GABA response^127^.

### Brain slice preparation and electrophysiology

All work on animals was carried out in accordance with the UK Animals (Scientific Procedures) Act 1986 under project and personal licenses granted by the UK Home Office. 250 μm thick transverse acute brain slices were prepared from adult (P67-84) male C57BL/6J mice using a Leica VT1200S vibroslicer. After terminal anaesthesia with isoflurane the brain was rapidly removed and sliced in an ice-cold slicing solution composed of (mM): 85 NaCl, 2.5 KCl, 1 CaCl_2_, 4 MgCl_2_, 1.25 NaH_2_PO_4_, 26 NaHCO_3_, 75 sucrose, and 25 glucose, 2 kynurenic acid, pH - 7.4; bubbled with 95% O_2_ and 5% CO_2_. The slicing solution was exchanged at 37 °C for 60 min under gravity flow with a recording solution or artificial cerebrospinal fluid (ACSF) containing (mM): 125 NaCl, 2.5 KCl, 2 CaCl_2_, 1 MgCl_2_, 1.25 NaH_2_PO_4_, 26 NaHCO_3_, 2 kynurenic acid, and 25 glucose, pH - 7.4.

Spontaneous inhibitory postsynaptic currents (sIPSCs) were recorded from hippocampal dentate gyrus granule cells (DGGCs) using patch electrodes of 4-5 MΩ resistance filled with an internal solution containing (mM): 120 CsCl 1 MgCl_2_, 11 EGTA, 30 KOH, 10 HEPES, 1 CaCl_2_, and 2 K_2_ATP; pH - 7.2. sIPSCs were recorded at −60 mV using 5 kHz filtering and optimal series resistance and whole-cell capacitance compensation.

sIPSCs were analysed using WinEDR and WinWCP (John Dempster, University of Strathclyde, UK). Percentage change in IPSC amplitudes, frequencies, rise times, decay times and charge transfers were calculated in the presence of nanobody or diazepam in comparison to controls in normal recording solution (aCSF). The statistical significance of sIPSC parameters in the presence or absence of treatments in each cell was assessed using a paired t-test in Graphpad Instat.

### Natural genetic variation in ligand-binding residues

To investigate the extent of natural genetic variation in the BZD binding site, ligand-binding residues were identified using inhouse written Perl scripts, available upon request from authors. Residues making atomic contacts with the ligands within 5Å distance were classified as ligand-binding residues. The number of such contacts were also calculated and are provided in the asteroid plots^128^. Conservation profiles for ligand binding positions across all human GBRA and GBRG subunits were then generated. We retrieved genetic variation data in different GBRA and GBRG receptors in 138,632 unrelated healthy humans from gnomAD database^54^. Genetic variation data was retrieved only for non-engineered positions and those that have identical residues as that of GBRA5 or GBRG2.

## REFERENCES

1 Morales-Perez, C. L., Noviello, C. M. & Hibbs, R. E. X-ray structure of the human alpha4beta2 nicotinic receptor. Nature 538, 411–415, doi:10.1038/nature19785 (2016).

2 Miyazawa, A., Fujiyoshi, Y. & Unwin, N. Structure and gating mechanism of the acetylcholine receptor pore. Nature 423, 949–955, doi:10.1038/nature01748 (2003).

3 Unwin, N. Refined structure of the nicotinic acetylcholine receptor at 4A resolution. Journal of molecular biology 346, 967–989, doi:10.1016/j.jmb.2004.12.031 (2005).

4 Hassaine, G. et al. X-ray structure of the mouse serotonin 5-HT3 receptor. Nature 512, 276–281, doi:10.1038/nature13552 (2014).

5 Miller, P. S. & Aricescu, A. R. Crystal structure of a human GABAA receptor. Nature 512, 270–275, doi:10.1038/nature13293 (2014).

6 Du, J., Lu, W., Wu, S., Cheng, Y. & Gouaux, E. Glycine receptor mechanism elucidated by electron cryo-microscopy. Nature 526, 224–229, doi:10.1038/nature14853 (2015).

7 Huang, X., Chen, H., Michelsen, K., Schneider, S. & Shaffer, P. L. Crystal structure of human glycine receptor-alpha3 bound to antagonist strychnine. Nature 526, 277–280, doi:10.1038/nature14972 (2015).

8 Hibbs, R. E. & Gouaux, E. Principles of activation and permeation in an anion-selective Cys-loop receptor. Nature 474, 54–60, doi:10.1038/nature10139 (2011).

9 Althoff, T., Hibbs, R. E., Banerjee, S. & Gouaux, E. X-ray structures of GluCl in apo states reveal a gating mechanism of Cys-loop receptors. Nature 512, 333–337, doi:10.1038/nature13669 (2014).

10 Nemecz, A., Prevost, M. S., Menny, A. & Corringer, P. J. Emerging Molecular Mechanisms of Signal Transduction in Pentameric Ligand-Gated Ion Channels. Neuron 90, 452–470, doi:10.1016/j.neuron.2016.03.032 (2016).

11 Olsen, R. W. & Sieghart, W. International Union of Pharmacology. LXX. Subtypes of gamma-aminobutyric acid(A) receptors: classification on the basis of subunit composition, pharmacology, and function. Update. Pharmacological reviews 60, 243–260, doi:10.1124/pr.108.00505 (2008).

12 Fritschy, J. M. & Panzanelli, P. GABAA receptors and plasticity of inhibitory neurotransmission in the central nervous system. The European journal of neuroscience 39, 1845–1865, doi:10.1111/ejn.12534 (2014).

13 Chang, Y., Wang, R., Barot, S. & Weiss, D. S. Stoichiometry of a recombinant GABAA receptor. The Journal of neuroscience: the official journal of the Society for Neuroscience 16, 5415–5424 (1996).

14 Collinson, N. et al. Enhanced learning and memory and altered GABAergic synaptic transmission in mice lacking the alpha 5 subunit of the GABAA receptor. The Journal of neuroscience: the official journal of the Society for Neuroscience 22, 5572–5580, doi:20026436 (2002).

15 Fernandez, F. et al. Pharmacotherapy for cognitive impairment in a mouse model of Down syndrome. Nature neuroscience 10, 411–413, doi:10.1038/nn1860 (2007).

16 Han, S. et al. Autistic-like behaviour in Scn1a+/−mice and rescue by enhanced GABA-mediated neurotransmission. Nature 489, 385–390, doi:10.1038/nature11356 (2012).

17 Clarkson, A. N., Huang, B. S., Macisaac, S. E., Mody, I. & Carmichael, S. T. Reducing excessive GABA-mediated tonic inhibition promotes functional recovery after stroke. Nature 468, 305–309, doi:10.1038/nature09511 (2010).

18 Yip, G. M. et al. A propofol binding site on mammalian GABAA receptors identified by photolabeling. Nature chemical biology 9, 715–720, doi:10.1038/nchembio.1340 (2013).

19 Jayakar, S. S. et al. Multiple propofol-binding sites in a gamma-aminobutyric acid type A receptor (GABAAR) identified using a photoreactive propofol analog. The Journal of biological chemistry 289, 27456–27468, doi:10.1074/jbc.M114.581728 (2014).

20 Sieghart, W., Ramerstorfer, J., Sarto-Jackson, I., Varagic, Z. & Ernst, M. A novel GABA(A) receptor pharmacology: drugs interacting with the alpha(+) beta(-) interface. British journal of pharmacology 166, 476–485, doi:10.1111/j.1476-5381.2011.01779.x (2012).

21 Laverty, D. et al. Crystal structures of a GABAA-receptor chimera reveal new endogenous neurosteroid-binding sites. Nature structural & molecular biology 24, 977–985, doi:10.1038/nsmb.3477 (2017).

22 Hanchar, H. J. et al. Ethanol potently and competitively inhibits binding of the alcohol antagonist Ro15-4513 to alpha4/6beta3delta GABAA receptors. Proceedings of the National Academy of Sciences of the United States of America 103, 8546–8551, doi:10.1073/pnas.0509903103 (2006).

23 Zeller, A., Arras, M., Jurd, R. & Rudolph, U. Identification of a molecular target mediating the general anesthetic actions of pentobarbital. Molecular pharmacology 71, 852–859, doi:10.1124/mol.106.030049 (2007).

24 Rudolph, U. & Knoflach, F. Beyond classical benzodiazepines: novel therapeutic potential of GABAA receptor subtypes. Nature reviews. Drug discovery 10, 685–697, doi:10.1038/nrd3502 (2011).

25 Li, G. D. et al. Identification of a GABAA receptor anesthetic binding site at subunit interfaces by photolabeling with an etomidate analog. The Journal of neuroscience: the official journal of the Society for Neuroscience 26, 11599–11605, doi:10.1523/JNEUROSCI.3467-06.2006 (2006).

26 Baldwin, D. S. et al. Benzodiazepines: risks and benefits. A reconsideration. Journal of psychopharmacology 27, 967–971, doi:10.1177/0269881113503509 (2013).

27 Knabl, J. et al. Reversal of pathological pain through specific spinal GABAA receptor subtypes. Nature 451, 330–334, doi:10.1038/nature06493 (2008).

28 Charych, E. I., Liu, F., Moss, S. J. & Brandon, N. J. GABA(A) receptors and their associated proteins: implications in the etiology and treatment of schizophrenia and related disorders. Neuropharmacology 57, 481–495, doi:10.1016/j.neuropharm.2009.07.027 (2009).

29 Sigel, E. & Steinmann, M. E. Structure, function, and modulation of GABA(A) receptors. The Journal of biological chemistry 287, 40224–40231, doi:10.1074/jbc.R112.386664 (2012).

30 Pritchett, D. B. et al. Importance of a novel GABAA receptor subunit for benzodiazepine pharmacology. Nature 338, 582–585, doi:10.1038/338582a0 (1989).

31 Pritchett, D. B., Luddens, H. & Seeburg, P. H. Type I and type II GABAA-benzodiazepine receptors produced in transfected cells. Science 245, 1389–1392 (1989).

32 Wieland, H. A., Luddens, H. & Seeburg, P. H. A single histidine in GABAA receptors is essential for benzodiazepine agonist binding. J Biol Chem 267, 1426–1429 (1992).

33 Amin, J., Brooks-Kayal, A. & Weiss, D. S. Two tyrosine residues on the alpha subunit are crucial for benzodiazepine binding and allosteric modulation of gamma-aminobutyric acidA receptors. MolPharmacol 51, 833–841 (1997).

34 Buhr, A., Schaerer, M. T., Baur, R. & Sigel, E. Residues at positions 206 and 209 of the alpha1 subunit of gamma-aminobutyric AcidA receptors influence affinities for benzodiazepine binding site ligands. Mol Pharmacol 52, 676–682 (1997).

35 Buhr, A. & Sigel, E. A point mutation in the gamma2 subunit of gamma-aminobutyric acid type A receptors results in altered benzodiazepine binding site specificity. Proceedings of the National Academy of Sciences of the United States of America 94, 8824–8829 (1997).

36 Buhr, A., Baur, R. & Sigel, E. Subtle changes in residue 77 of the gamma subunit of alpha1beta2gamma2 GABAA receptors drastically alter the affinity for ligands of the benzodiazepine binding site. The Journal of biological chemistry 272, 11799–11804 (1997).

37 Kucken, A. M. et al. Identification of benzodiazepine binding site residues in the gamma2 subunit of the gamma-aminobutyric acid(A) receptor. Molecular pharmacology 57, 932–939 (2000).

38 Hunkeler, W. et al. Selective antagonists of benzodiazepines. Nature 290, 514–516 (1981).

39 Puia, G., Ducic, I., Vicini, S. & Costa, E. Molecular mechanisms of the partial allosteric modulatory effects of bretazenil at gamma-aminobutyric acid type A receptor. Proceedings of the National Academy of Sciences of the United States of America 89, 3620–3624 (1992).

40 Jansen, M., Bali, M. & Akabas, M. H. Modular design of Cys-loop ligand-gated ion channels: functional 5-HT3 and GABA rho1 receptors lacking the large cytoplasmic M3M4 loop. The Journal of general physiology 131, 137–146, doi:10.1085/jgp.200709896 (2008).

41 Chu, R. et al. Redesign of a four-helix bundle protein by phage display coupled with proteolysis and structural characterization by NMR and X-ray crystallography. Journal of molecular biology 323, 253–262 (2002).

42 Baumann, S. W., Baur, R. & Sigel, E. Forced subunit assembly in alpha1beta2gamma2 GABAA receptors. Insight into the absolute arrangement. The Journal of biological chemistry 277, 46020–46025, doi:10.1074/jbc.M207663200 (2002).

43 Tretter, V., Ehya, N., Fuchs, K. & Sieghart, W. Stoichiometry and assembly of a recombinant GABAA receptor subtype. The Journal of neuroscience: the official journal of the Society for Neuroscience 17, 2728–2737 (1997).

44 Luddens, H. et al. Cerebellar GABAA receptor selective for a behavioural alcohol antagonist. Nature 346, 648–651, doi:10.1038/346648a0 (1990).

45 Wong, G. et al. Synthetic and computer-assisted analysis of the structural requirements for selective, high-affinity ligand binding to diazepam-insensitive benzodiazepine receptors. J Med Chem 36, 1820–1830 (1993).

46 Sigel, E., Schaerer, M. T., Buhr, A. & Baur, R. The benzodiazepine binding pocket of recombinant alpha1beta2gamma2 gamma-aminobutyric acidA receptors: relative orientation of ligands and amino acid side chains. Molecular pharmacology 54, 1097–1105 (1998).

47 Hunkeler, W. Benzodiazepines, the Story of the Antagonist Flumazenil and of the Partial Agonist Bretazenil. Chimia 47, 141–147 (1993).

48 Wallner, M., Hanchar, H. J. & Olsen, R. W. Low-dose alcohol actions on alpha4beta3delta GABAA receptors are reversed by the behavioral alcohol antagonist Ro15-4513. Proceedings of the National Academy of Sciences of the United States of America 103, 8540–8545, doi:10.1073/pnas.0600194103 (2006).

49 Mendez, M. A. et al. The brain GABA-benzodiazepine receptor alpha-5 subtype in autism spectrum disorder: a pilot [(11)C]Ro15-4513 positron emission tomography study. Neuropharmacology 68, 195–201, doi:10.1016/j.neuropharm.2012.04.008 (2013).

50 Sawyer, G. W., Chiara, D. C., Olsen, R. W. & Cohen, J. B. Identification of the bovine gamma-aminobutyric acid type A receptor alpha subunit residues photolabeled by the imidazobenzodiazepine [3H]Ro15-4513. The Journal of biological chemistry 277, 50036–50045, doi:10.1074/jbc.M209281200 (2002).

51 Kucken, A. M., Teissere, J. A., Seffinga-Clark, J., Wagner, D. A. & Czajkowski, C. Structural requirements for imidazobenzodiazepine binding to GABA(A) receptors. Molecular pharmacology 63, 289–296 (2003).

52 Ananthan, S. et al. Synthesis and structure-activity relationships of 3,5-disubstituted 4,5-dihydro-6H-imidazo[1,5-a][1,4]benzodiazepin-6-ones at diazepam-sensitive and diazepam-insensitive benzodiazepine receptors. J Med Chem 36, 479–490 (1993).

53 Watjen, F. et al. Novel Benzodiazepine Receptor Partial Agonists - Oxadiazolylimidazobenzodiazepines. Journal of Medicinal Chemistry 32, 2282–2291, doi:Doi 10.1021/Jm00130a010 (1989).

54 Lek, M. et al. Analysis of protein-coding genetic variation in 60,706 humans. Nature 536, 285–291, doi:10.1038/nature19057 (2016).

55 Newell, J. G., McDevitt, R. A. & Czajkowski, C. Mutation of glutamate 155 of the GABAA receptor beta2 subunit produces a spontaneously open channel: a trigger for channel activation. The Journal of neuroscience: the official journal of the Society for Neuroscience 24, 11226–11235, doi:10.1523/JNEUROSCI.3746-04.2004 (2004).

56 Dunn, S. M., Davies, M., Muntoni, A. L. & Lambert, J. J. Mutagenesis of the rat alpha1 subunit of the gamma-aminobutyric acid(A) receptor reveals the importance of residue 101 in determining the allosteric effects of benzodiazepine site ligands. Molecular pharmacology 56, 768–774 (1999).

57 Hanson, S. M., Morlock, E. V., Satyshur, K. A. & Czajkowski, C. Structural requirements for eszopiclone and zolpidem binding to the gamma-aminobutyric acid type-A (GABAA) receptor are different. J Med Chem 51, 7243–7252, doi:10.1021/jm800889m (2008).

58 Rudolph, U. et al. Benzodiazepine actions mediated by specific gamma-aminobutyric acid(A) receptor subtypes. Nature 401, 796–800, doi:10.1038/44579 (1999).

59 Pirker, S., Schwarzer, C., Wieselthaler, A., Sieghart, W. & Sperk, G. GABA(A) receptors: immunocytochemical distribution of 13 subunits in the adult rat brain. Neuroscience 101, 815–850 (2000).

60 Aebi, M. N-linked protein glycosylation in the ER. Biochimica et biophysica acta 1833, 2430–2437, doi:10.1016/j.bbamcr.2013.04.001 (2013).

61 Calimet, N. et al. A gating mechanism of pentameric ligand-gated ion channels. Proceedings of the National Academy of Sciences of the United States of America 110, E3987–3996, doi:10.1073/pnas.1313785110 (2013).

62 Miller, P. S. et al. Structural basis for GABAA receptor potentiation by neurosteroids. Nat StructMolBiol 24, 986–992, doi:10.1038/nsmb.3484 (2017).

63 Miller, P. S. & Smart, T. G. Binding, activation and modulation of Cys-loop receptors. Trends in pharmacological sciences 31, 161–174, doi:10.1016/j.tips.2009.12.005 (2010).

64 Hansen, S. B., Wang, H. L., Taylor, P. & Sine, S. M. An ion selectivity filter in the extracellular domain of Cys-loop receptors reveals determinants for ion conductance. The Journal of biological chemistry 283, 36066–36070, doi:10.1074/jbc.C800194200 (2008).

65 Moroni, M., Meyer, J. O., Lahmann, C. & Sivilotti, L. G. In glycine and GABA(A) channels, different subunits contribute asymmetrically to channel conductance via residues in the extracellular domain. The Journal of biological chemistry 286, 13414–13422, doi:10.1074/jbc.M110.204610 (2011).

66 Mancinelli, R., Botti, A., Bruni, F., Ricci, M. A. & Soper, A. K. Hydration of sodium, potassium, and chloride ions in solution and the concept of structure maker/breaker. The journal of physical chemistry. B 111, 13570–13577, doi:10.1021/jp075913v (2007).

67 Gielen, M., Thomas, P. & Smart, T. G. The desensitization gate of inhibitory Cys-loop receptors. Nature communications 6, 6829, doi:10.1038/ncomms7829 (2015).

68 Fatima-Shad, K. & Barry, P. H. Anion permeation in GABA- and glycine-gated channels of mammalian cultured hippocampal neurons. Proceedings. Biological sciences 253, 69–75, doi:10.1098/rspb.1993.0083 (1993).

69 Rao, S., Klesse, G., Stansfeld, P. J., Tucker, S. J. & Sansom, M. S. P. A BEST example of channel structure annotation by molecular simulation. Channels 11, 347–353, doi:10.1080/19336950.2017.1306163 (2017).

70 Trick, J. L. et al. Functional Annotation of Ion Channel Structures by Molecular Simulation. Structure 24, 2207–2216, doi:10.1016/j.str.2016.10.005 (2016).

71 Mortensen, M. & Smart, T. G. Extrasynaptic alphabeta subunit GABAA receptors on rat hippocampal pyramidal neurons. The Journal of physiology 577, 841–856, doi:10.1113/jphysiol.2006.117952 (2006).

72 Luu, T., Cromer, B., Gage, P. W. & Tierney, M. L. A role for the 2’ residue in the second transmembrane helix of the GABA A receptor gamma2S subunit in channel conductance and gating. The Journal of membrane biology 205, 17–28, doi:10.1007/s00232-005-0759-2 (2005).

73 Beato, M., Groot-Kormelink, P. J., Colquhoun, D. & Sivilotti, L. G. Openings of the rat recombinant alpha 1 homomeric glycine receptor as a function of the number of agonist molecules bound. The Journal of general physiology 119, 443–466 (2002).

74 Mortensen, M. et al. Activation of single heteromeric GABA(A) receptor ion channels by full and partial agonists. The Journal of physiology 557, 389–413, doi:10.1113/jphysiol.2003.054734 (2004).

75 Unwin, N. & Fujiyoshi, Y. Gating movement of acetylcholine receptor caught by plunge-freezing. Journal of molecular biology 422, 617–634, doi:10.1016/j.jmb.2012.07.010 (2012).

76 Aricescu, A. R., Lu, W. & Jones, E. Y. A time- and cost-efficient system for high-level protein production in mammalian cells. Acta crystallographica. Section D, Biological crystallography 62, 1243–1250, doi:10.1107/S0907444906029799 (2006).

77 Molday, R. S. & MacKenzie, D. Monoclonal antibodies to rhodopsin: characterization, cross-reactivity, and application as structural probes. Biochemistry 22, 653–660 (1983).

78 Oprian, D. D., Molday, R. S., Kaufman, R. J. & Khorana, H. G. Expression of a synthetic bovine rhodopsin gene in monkey kidney cells. Proceedings of the National Academy of Sciences of the United States of America 84, 8874–8878 (1987).

79 Taylor, P. M. et al. Identification of amino acid residues within GABA(A) receptor beta subunits that mediate both homomeric and heteromeric receptor expression. The Journal of neuroscience: the official journal of the Society for Neuroscience 19, 6360–6371 (1999).

80 Reeves, P. J., Callewaert, N., Contreras, R. & Khorana, H. G. Structure and function in rhodopsin: high-level expression of rhodopsin with restricted and homogeneous N-glycosylation by a tetracycline-inducible N-acetylglucosaminyltransferase I-negative HEK293S stable mammalian cell line. Proceedings of the National Academy of Sciences of the United States of America 99, 13419–13424, doi:10.1073/pnas.212519299 (2002).

81 Aricescu, A. R. & Owens, R. J. Expression of recombinant glycoproteins in mammalian cells: towards an integrative approach to structural biology. Current opinion in structural biology 23, 345–356, doi:10.1016/j.sbi.2013.04.003 (2013).

82 Zacharias, D. A., Violin, J. D., Newton, A. C. & Tsien, R. Y. Partitioning of lipid-modified monomeric GFPs into membrane microdomains of live cells. Science 296, 913–916, doi:10.1126/science.1068539 (2002).

83 Nagai, T. et al. A variant of yellow fluorescent protein with fast and efficient maturation for cell-biological applications. Nature biotechnology 20, 87–90, doi:10.1038/nbt0102-87 (2002).

84 Pardon, E. et al. A general protocol for the generation of Nanobodies for structural biology. Nature protocols 9, 674–693, doi:10.1038/nprot.2014.039 (2014).

85 Chang, V. T. et al. Glycoprotein structural genomics: solving the glycosylation problem. Structure 15, 267–273, doi:10.1016/j.str.2007.01.011 (2007).

86 Walter, T. S. et al. A procedure for setting up high-throughput nanolitre crystallization experiments. Crystallization workflow for initial screening, automated storage, imaging and optimization. Acta crystallographica. Section D, Biological crystallography 61, 651–657, doi:10.1107/S0907444905007808 (2005).

87 Parker, J. L. & Newstead, S. Current trends in alpha-helical membrane protein crystallization: an update. Protein science: a publication of the Protein Society 21, 1358–1365, doi:10.1002/pro.2122 (2012).

88 Otwinowski, Z. & Minor, W. Processing of X-ray Diffraction Data Collected in Oscillation Mode. Methods in enzymology 276, 307–326, doi:10.1016/S0076-6879(97)76066-X (1997).

89 Tickle, i. J. et al. STARANISO. Global Phasing Ltd., Cambridge, United Kingdom. (2018).

90 Winn, M. D. et al. Overview of the CCP4 suite and current developments. Acta crystallographica. Section D, Biological crystallography 67, 235–242, doi:10.1107/S0907444910045749 (2011).

91 Evans, P. R. & Murshudov, G. N. How good are my data and what is the resolution? Acta crystallographica. Section D, Biological crystallography 69, 1204–1214, doi:10.1107/S0907444913000061 (2013).

92 Vonrhein, C. et al. Data processing and analysis with the autoPROC toolbox. Acta crystallographica. Section D, Biological crystallography 67, 293–302, doi:10.1107/S0907444911007773 (2011).

93 McCoy, A. J. et al. Phaser crystallographic software. Journal of Applied Crystallography 40, 658–674, doi:10.1107/S0021889807021206 (2007).

94 Emsley, P., Lohkamp, B., Scott, W. G. & Cowtan, K. Features and development of Coot. Acta crystallographica. Section D, Biological crystallography 66, 486–501, doi:10.1107/S0907444910007493 (2010).

95 Bricogne, G. et al. BUSTER version X.Y.Z., Global Phasing Ltd., Cambridge, United Kingdom. (2017).

96 Murshudov, G. N. et al. REFMAC5 for the refinement of macromolecular crystal structures. Acta crystallographica. Section D, Biological crystallography 67, 355–367, doi:10.1107/S0907444911001314 (2011).

97 Adams, P. D. et al. PHENIX: a comprehensive Python-based system for macromolecular structure solution. Acta crystallographica. Section D, Biological crystallography 66, 213–221, doi:10.1107/S0907444909052925 (2010).

98 Afonine, P. V. et al. Towards automated crystallographic structure refinement with phenix.refine. Acta crystallographica. Section D, Biological crystallography 68, 352–367, doi:10.1107/S0907444912001308 (2012).

99 Smart, O. S. et al. grade, v1.105, Global Phasing Ltd., Cambridge, United Kingdom. (2011).

100 Chen, V. B. et al. MolProbity: all-atom structure validation for macromolecular crystallography. Acta crystallographica. Section D, Biological crystallography 66, 12–21, doi:10.1107/S0907444909042073 (2010).

101 Pettersen, E. F. et al. UCSF Chimera--a visualization system for exploratory research and analysis. J Comput Chem 25, 1605–1612, doi:10.1002/jcc.20084 (2004).

102 Li, X. et al. Electron counting and beam-induced motion correction enable near-atomic-resolution single-particle cryo-EM. Nature methods 10, 584–590, doi:10.1038/nmeth.2472 (2013).

103 Rohou, A. & Grigorieff, N. CTFFIND4: Fast and accurate defocus estimation from electron micrographs. Journal of structural biology 192, 216–221, doi:10.1016/j.jsb.2015.08.008 (2015).

104 Scheres, S. H. RELION: implementation of a Bayesian approach to cryo-EM structure determination. Journal of structural biology 180, 519–530, doi:10.1016/j.jsb.2012.09.006 (2012).

105 Tang, G. et al. EMAN2: an extensible image processing suite for electron microscopy. Journal of structural biology 157, 38–46, doi:10.1016/j.jsb.2006.05.009 (2007).

106 Zheng, S. Q. et al. MotionCor2: anisotropic correction of beam-induced motion for improved cryo-electron microscopy. Nature methods 14, 331–332, doi:10.1038/nmeth.4193 (2017).

107 Terwilliger, T. C., Sobolev, O., Afonine, P. V. & Adams, P. D. Automated map sharpening by maximization of detail and connectivity. bioRxiv-Acta Crystallographica Section D (2018).

108 Song, Y. et al. High-resolution comparative modeling with RosettaCM. Structure 21, 1735–1742, doi:10.1016/j.str.2013.08.005 (2013).

109 Afonine, P. V., Headd, J. J., Terwilliger, T. C. & Adams, P. D. New tool: phenix.real_space_refine. Computational Crystallography Newsletter 4, 43–44 (2013).

110 Davis, I. W. et al. MolProbity: all-atom contacts and structure validation for proteins and nucleic acids. Nucleic acids research 35, W375–383, doi:10.1093/nar/gkm216 (2007).

111 Stuart, D. I., Levine, M., Muirhead, H. & Stammers, D. K. Crystal structure of cat muscle pyruvate kinase at a resolution of 2.6 A. Journal of molecular biology 134, 109–142 (1979).

112 Smart, O. S., Neduvelil, J. G., Wang, X., Wallace, B. A. & Sansom, M. S. HOLE: a program for the analysis of the pore dimensions of ion channel structural models. J Mol Graph 14, 354–360, 376 (1996).

113 Pavelka, A. et al. CAVER: Algorithms for Analyzing Dynamics of Tunnels in Macromolecules. IEEE/ACM transactions on computational biology and bioinformatics 13, 505–517, doi:10.1109/TCBB.2015.2459680 (2016).

114 Trott, O. & Olson, A. J. AutoDock Vina: improving the speed and accuracy of docking with a new scoring function, efficient optimization, and multithreading. J Comput Chem 31, 455–461, doi:10.1002/jcc.21334 (2010).

115 Smart, O. S. et al. Grade version 1.104. Cambridge, United Kingdom, Global Phasing Ltd. http://www.globalphasing.com (2011).

116 Stansfeld, P. J. & Sansom, M. S. Molecular simulation approaches to membrane proteins. Structure 19, 1562–1572, doi:10.1016/j.str.2011.10.002 (2011).

117 Jorgensen, W. L., Chandrasekhar, J., Madura, J. D., Impey, R. W. & Klein, M. L. Comparison of Simple Potential Functions for Simulating Liquid Water. J Chem Phys 79, 926–935, doi:Doi 10.1063/1.445869 (1983).

118 Van Der Spoel, D. et al. GROMACS: fast, flexible, and free. J Comput Chem 26, 1701–1718, doi:10.1002/jcc.20291 (2005).

119 Abraham, M. J. et al. GROMACS: high performance molecular simulations through multi-level parallelism from laptops to supercomputers. Software X 1–2, 19–25 (2015).

120 Jorgensen, W. L., Maxwell, D. S. & TiradoRives, J. Development and testing of the OPLS all-atom force field on conformational energetics and properties of organic liquids. Journal of the American Chemical Society 118, 11225–11236, doi:Doi 10.1021/Ja9621760 (1996).

121 Bussi, G., Donadio, D. & Parrinello, M. Canonical sampling through velocity rescaling. J Chem Phys 126, 014101, doi:10.1063/1.2408420 (2007).

122 Parrinello, M. & Rahman, A. Polymorphic Transitions in Single-Crystals - a New Molecular-Dynamics Method. JApplPhys 52, 7182–7190, doi:Doi 10.1063/1.328693 (1981).

123 Darden, T., York, D. & Pedersen, L. Particle Mesh Ewald - an N.Log(N) Method for Ewald Sums in Large Systems. J Chem Phys 98, 10089–10092, doi:Doi 10.1063/1.464397 (1993).

124 Hess, B., Bekker, H., Berendsen, H. J. C. & Fraaije, J. G. E. M. LINCS: A linear constraint solver for molecular simulations. Journal of Computational Chemistry 18, 1463–1472, doi:Doi 10.1002/(Sici)1096-987x(199709)18:12<1463::Aid-Jcc4>3.0.Co;2-H (1997).

125 Cheng, Y. & Prusoff, W. H. Relationship between the inhibition constant (K1) and the concentration of inhibitor which causes 50 per cent inhibition (I50) of an enzymatic reaction. Biochemical pharmacology 22, 3099–3108 (1973).

126 Hattori, M., Hibbs, R. E. & Gouaux, E. A fluorescence-detection size-exclusion chromatography-based thermostability assay for membrane protein precrystallization screening. Structure 20, 1293–1299, doi:10.1016/j.str.2012.06.009 (2012).

127 Hosie, A. M., Dunne, E. L., Harvey, R. J. & Smart, T. G. Zinc-mediated inhibition of GABA(A) receptors: discrete binding sites underlie subtype specificity. Nature neuroscience 6, 362–369, doi:10.1038/nn1030 (2003).

128 Kayikci, M. et al. Visualization and analysis of non-covalent contacts using the Protein Contacts Atlas. Nature structural & molecular biology 25, 185–194, doi:10.1038/s41594-017-0019-z (2018).

129 Baur, R., Luscher, B. P., Richter, L. & Sigel, E. A residue close to alpha1 loop F disrupts modulation of GABAA receptors by benzodiazepines while their binding is maintained. J Neurochem 115, 1478–1485, doi:10.1111/j.1471-4159.2010.07052.x (2010).

130 Rudolph, U. et al. Benzodiazepine actions mediated by specific gamma-aminobutyric acid(A) receptor subtypes. Nature 401, 796–800, doi:10.1038/44579 (1999).

